# Conserved auxin signalling synergism through the canonical and ETT-mediated pathways promotes gynoecium development in members of the *Brassicaceae*

**DOI:** 10.1101/2023.03.02.530771

**Authors:** Heather Marie McLaughlin, Tian-Feng Lü, Bhavani Natarajan, Lars Østergaard, Yang Dong

## Abstract

Gynoecium polarity establishment is regulated by auxin, a phytohormone whose signal is transduced through several pathways. The relationship between the ETT and canonical TIR1/AFB pathways, and their relevance for carpel development beyond *Arabidopsis thaliana*, have not been investigated. The data presented here show that the expression patterns of canonical and ETT-mediated signalling components, and phenotypes of higher order mutants are shared between *Arabidopsis* and *Capsella rubella*. *tir1 afb2 ett* mutants partially phenocopy *ett arf4* double mutants, suggesting a role for AUXIN RESPONSE FACTOR 4 (ARF4) in the integration of canonical and ETT-mediated signalling. Comparative transcriptomics revealed that the auxin-independent mis-regulation of *YABBY* genes correlate with patterning defects observed in *Arabidopsis ett arf4* mutants. Together, the data presented suggest conserved synergism between canonical and ETT-mediated pathways in gynoecium polarity establishment in the *Brassicaceae*. Finally, the data suggest that ETT/ARF4 function to prevent the auxin-induced expression of a range of targets in *Arabidopsis*, consistent with activator-repressor ARF antagonism, and implying that the maintenance of auxin insensitivity by repressive ARFs is important for a range of biological processes.

**Summary Statement:** Here the relationship between canonical and ETT-mediated auxin signalling machineries is investigated in *Arabidopsis* and *Capsella* revealing conserved synergism between these two pathways in *Brassicaceae* with distinct fruit shapes.

## Introduction

The gynoecium is the female reproductive organ produced in flowering plants which develops into the fruit. The gynoecium is produced in the inner-most whorl of the flower and is composed of several specialized tissues (Sessions and Zambryski, 1995). The gynoecium can be subdivided into three distinct regions along the apical-basal axis; the stigma, style; and ovary. The ovary is made up of two valves, which are separated from one another by the valve margins and medial replum. Gynoecium patterning and morphogenesis is dependent on the dynamic distribution of the phytohormone auxin (Moubayidin and Østergaard, 2014).

Auxin signalling occurs via several distinct pathways in *Arabidopsis* (Dharmasiri et al., 2005; Kepinski and Leyser, 2005; Simonini et al., 2016; Fendrych et al., 2018; Cao et al., 2019; Kuhn et al., 2020; Friml et al., 2022). The canonical auxin signalling pathway induces the expression of auxin sensitive genes in the nucleus (Paponov et al., 2008a), and is composed of three main components: Auxin/Indole-3-Acetic Acid repressor proteins (AUX/IAAs), Transport Inhibitor Response1/Arabidopsis F-box (TIR1/AFB) receptors, and Auxin Response Factors (ARFs). In the absence of auxin, ARFs are bound by AUX/IAAs, which recruit the TOPLESS (TPL) co-repressor, preventing the expression of auxin responsive genes (Szemenyei et al., 2008). Auxin increases the affinity of the TIR1/AFBs for AUX/IAAs. AUX/IAAs are subsequently degraded by the proteasome, allowing ARFs to promote auxin responsive gene expression (Dharmasiri et al., 2005; Kepinski and Leyser, 2005).

ARFs are classified into three distinct clades: A-ARFs, B-ARFs, and C-ARFs (Ulmasov et al., 1999; Tiwari et al., 2003; Finet et al., 2013). A-ARFs are transcriptional activators which readily interact with AUX/IAAs, whilst B and C ARFs are characterized as repressive, and interact with AUX/IAAs less readily (Ulmasov et al., 1999; Tiwari et al., 2003; Finet et al., 2013; Vernoux et al., 2011; reviewed in Guilfoyle & Hagen, 2012).

The ETTIN (ETT/ARF3)-mediated pathway was elucidated more recently in the context of its role in gynoecium development (Simonini et al., 2016; Kuhn et al., 2020). ETT is a B-class repressive ARF which lacks the C-terminal domain required for AUX/IAA interaction (Simonini et al., 2016), and functions independently of the canonical pathway (Kuhn et al., 2020). ETT recruits a TPL-HISTONE DEACETYLASE 19 (HDA19) complex to prevent the transcription of target loci in the absence of auxin. ETT interacts with auxin through its C-terminal ETTIN-SPECIFIC (ES) domain and undergoes a conformational change upon auxin binding which breaks the ETT-TPL-HDA19 complex. Histones then become acetylated, and genes are derepressed (Kuhn et al., 2020).

*ett* mutants exhibit gynoecium defects which phenocopy aberrations produced upon treatment with the auxin transport inhibitor N-1-Naphthylphthalamic Acid (NPA), highlighting the importance of ETT-mediated signalling for auxin related processes in carpel development (Nemhauser et al., 2000). Gynoecium morphogenesis is dependent on auxin in *Arabidopsis thaliana* (from here on, Arabidopsis; Moubayidin and Østergaard, 2014). However, studies on auxin signal transduction in the carpels have mainly focused on the role of the ETT-mediated pathway (Simonini et al., 2016; Simonini et al., 2017; Kuhn et al., 2020). An understanding of the relative contribution of canonical and ETT-mediated signalling to gynoecium development, and the relevance of this relationship outside of *Arabidopsis* has not been described.

Several canonical auxin receptors - *TIR1*, *AFB1*, *AFB2*, and *AFB3 -* are expressed in the style of the developing gynoecium (Dharmasiri et al., 2005; Cecchetti et al., 2008). Higher order *tir1afb1245* mutants exhibit curled siliques and reduced fertility indicative of gynoecium defects (Prigge et al., 2020). AUX/IAA-interacting ARFs (Vernoux et al., 2011; Piya et al., 2014), are also involved in the regulation of gynoecium development (Nagpal et al., 2005; Wang et al., 2005; Pekker et al., 2005; Kelley et al., 2012), further supporting a role for the canonical pathway in this process.

ETT shares partial redundancy with its paralogous ARF, ARF4, which contains a C-terminal PB1-domain and is capable of AUX/IAA interaction (Pekker et al., 2005; Kelley et al., 2012; Finet et al., 2010; Vernoux et al., 2011). However, whilst the transcriptional targets of ETT in the gynoecium are well described (Simonini et al., 2017; Andres-Robin et al., 2020), those of ARF4 are not, and little is known about the roles of ETT and ARF4 in gynoecium development beyond *Arabidopsis*.

The *Brassicaceae* family display an impressive array of fruit shape diversity including for example cylindrical, spherical, disc and heart-shaped fruits (Łangowski et al., 2016). Interestingly, gynoecia from species producing highly divergent fruit shapes still maintain the same general organization of tissues with two or more carpels and an apical radial style topped with stigmatic papillae (Langowski et al., 2016). Members of the *Brassicaceae* family therefore provide excellent models to study conserved mechanisms of carpel development. It was recently shown that gynoecium development in *Capsella rubella* (from here on *Capsella*), which diverged from *Arabidopsis* around 8 million years ago, and produces heart shaped fruits, is also dependent on auxin dynamics throughout development (Dong et al., 2019). However, the relative contributions of the canonical and ETT-mediated pathway to signal transduction in *Capsella* fruit development are not known.

Here, *tir1 afb2 ett* mutants were generated in *Arabidopsis and Capsella.* In both species, the triple mutant showed exacerbation of the *ett* phenotype suggesting that the canonical and ETT-mediated pathways function synergistically in gynoecium development, and that this is conserved within the *Brassicaceae*. Furthermore, *the tir1 afb2 ett* triple mutant partially phenocopied the *ett arf4* double mutant in both species, suggesting a role for ARF4 in the integration of signalling between the canonical and ETT-mediated pathways. A comparative transcriptomics analysis was conducted to identify distinct and overlapping ETT and ARF4 targets in *Arabidopsis*. The data presented suggest that ETT and ARF4 regulate distinct and overlapping targets independently of auxin including YABBY transcription factors (TFs), and the mis-regulation of these genes correlates with the phenotypes observed in the *Arabidopsis* mutants.

Finally, auxin sensitive transcriptomics in *Arabidopsis* suggest that ETT and ARF4 maintain the auxin insensitivity of a range of target loci, many which became upregulated in response to auxin upon their mutation, suggesting that A-B ARF antagonism previously observed in *Marchantia* (Kato et al., 2020) and *Physcomitrium* (Lavy et al., 2016) may regulate a range of biological processes in *Arabidopsis*.

Together, the findings presented reveal conserved synergism between the canonical and ETT-mediated pathway machineries in *Arabidopsis* and *Capsella*, identify auxin independent targets of ETT and ARF4 in *Arabidopsis*, and suggest that the maintenance of auxin insensitivity at target loci plays an important role in the regulation of a range of responses to environmental stresses.

## Results

### Canonical and ETT-mediated pathway components are co-expressed during gynoecium development in *Arabidopsis* and *Capsella*

To assess whether the canonical and ETT-mediated pathways may co-regulate processes during *Arabidopsis* gynoecium, we first studied the expression of GUS reporter lines for *ETT*, *TIR1*, *AFB2* and *AFB3* during gynoecium development (Fig. 1). In *Arabidopsis*, *ETT* is expressed throughout the valves of the gynoecium, and is most highly expressed at the apex of the valves in the region which gives rise to the style at stage 10-12 (Fig. 1). At later stages *ETT* expression is much lower in the valves and becomes mainly restricted to the style and replum. *AtTIR1*, *AtAFB2* and *AtAFB3* are also expressed throughout the valves during stage 10-11 of gynoecium development, like *AtETT* (Fig. 1). *AtTIR1* expression becomes reduced in the valves after stage 13 of development and is restricted to the style and replum at stage 14. *AtAFB2* is also restricted to the style from stage 13 of gynoecium development, whilst *AtAFB3* expression remains strongest in the apical valves, stigma, and style at stage 12-13 (Fig. 1). *AtAFB3* is then restricted to the style and stigma from stage 14 of gynoecium development.

**Fig. 1:**
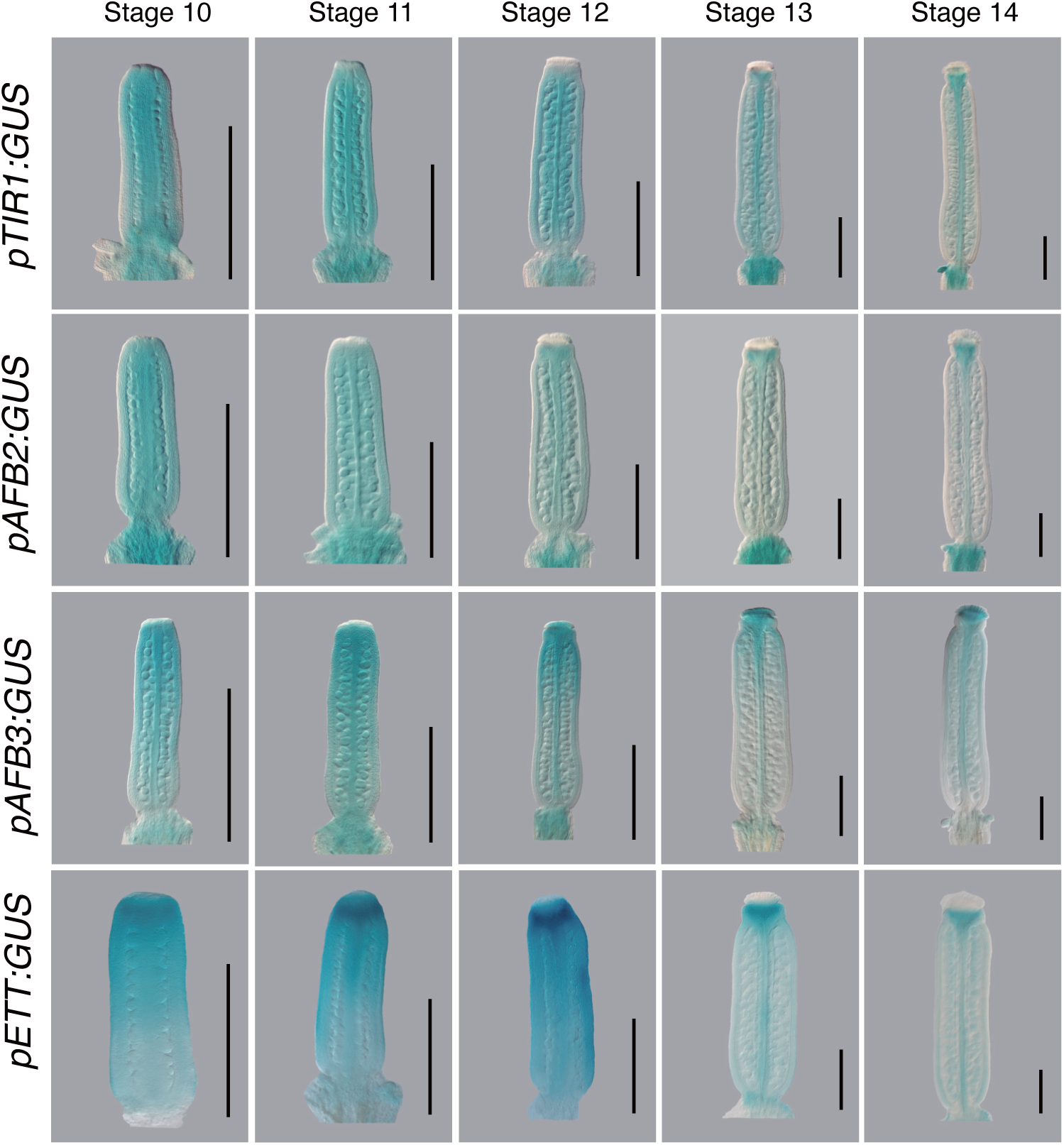
Expression analysis of genes involved in the canonical and ETT-mediated auxin signalling pathway in *Arabidopsis thaliana.* GUS staining showing the expression patterns of *pAtETT, pAtTIR1*, *pAtAFB2* and *pAtAFB3* from stage 10-14 of gynoecium development. Scale bars 500μm.

The expression of the canonical and ETT-mediated signalling machinery within the same domains suggests that both pathways may be involved in gynoecium development, particularly in valve and style development in *Arabidopsis*.

To assess whether a role for both pathways is conserved, similar reporter lines were generated to assess the expression patterns of *ETT*, *TIR1*, *AFB2* and *AFB3* in *Capsella* (Fig. 2). *CrETT* is expressed strongly in the apical tissues of the gynoecium and the ovules during stage 10*, whilst CrTIR1, CrAFB2, and CrAFB3* are expressed ubiquitously across the gynoecium at this stage (Fig. 2). *CrETT* is then most strongly expressed in the style from stage 11-14. *CrTIR1*, *CrAFB2* and *CrAFB3* are continually expressed in the apical valves and style from stage 11-13, and at stage 14 their expression levels are highest in the outgrowing shoulders of the *Capsella* fruit, although *CrTIR1* is expressed at lower levels than *CrAFB2* and *CrAFB3* (Fig. 2).

**Fig. 2:**
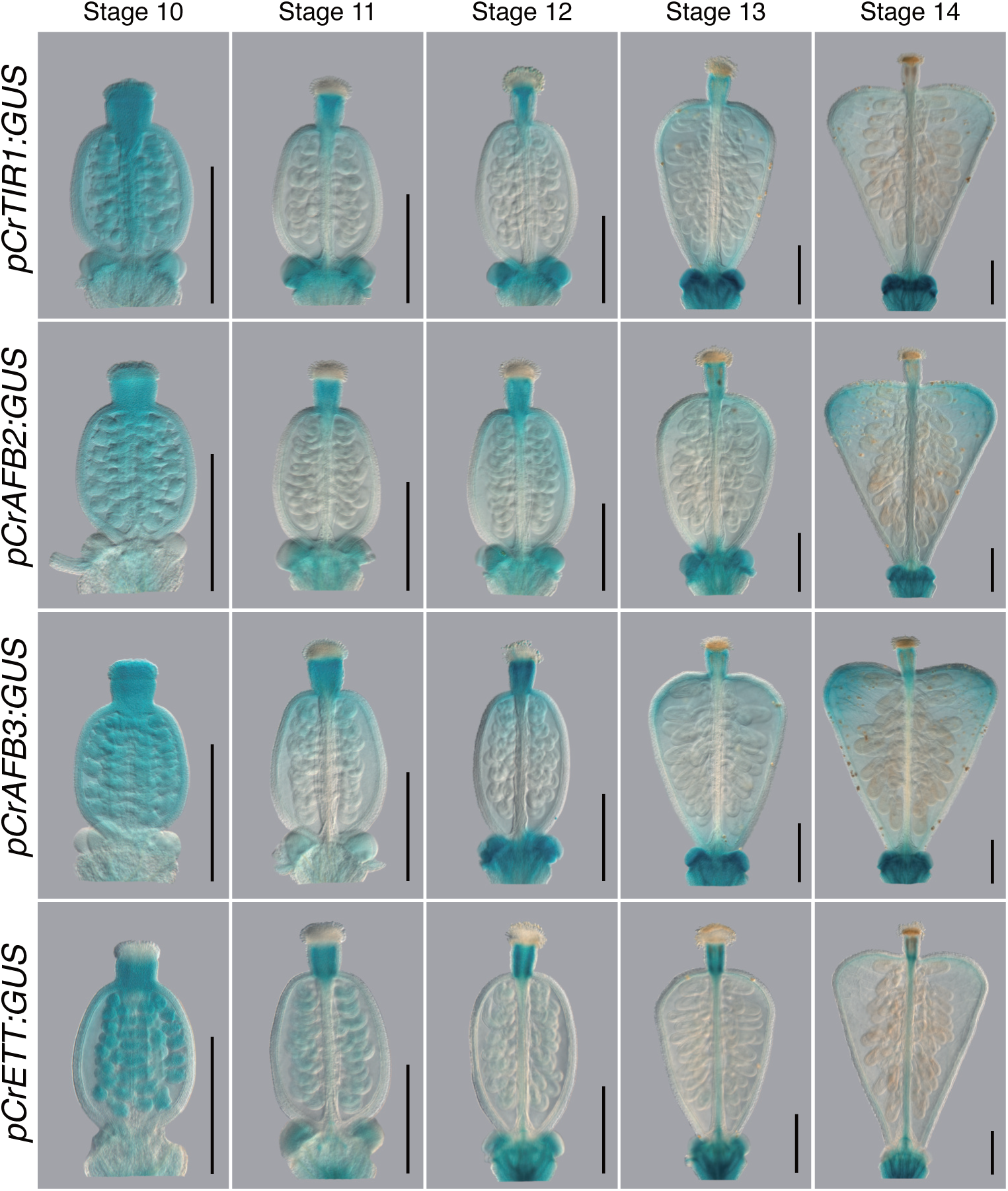
Expression analysis of genes involved in the canonical and ETT-mediated auxin signalling pathway in *Capsella rubella*. GUS staining showing the expression patterns of *pCrETT, pCrTIR1*, *pCrAFB2* and *pCrAFB3* from stage 10-14 of gynoecium development. Scale bars 500μm.

Together, these data suggest shared roles for the canonical and ETT-mediated pathways in valve and style development in both *Arabidopsis* and *Capsella* and suggest a role for the canonical pathway in post fertilization valve outgrowth in *Capsella*, which may contribute to heart-shaped fruit formation.

### Phenotypic analyses suggest synergism between canonical and ETT-mediated pathways is conserved in other members of the *Brassicaceae*

To understand the relationship between the canonical and ETT-mediated auxin signalling pathways during gynoecium development, *tir1-1 afb2-3 ett-3* triple mutants were generated in *Arabidopsis*, and the resulting phenotypes were assessed compared to single and double mutants (Fig. 3). ETT functions partly redundantly with ARF4, which shares ETT’s expression domain in the valves and apical tissues of the gynoecium during development (Fig. S1). ARF4 interacts with the canonical pathway AUX/IAAs (Vernoux et al., 2011), implicating it in the canonical pathway. The *arf4-2* single mutants and *ett-3 arf4-2* double mutants were therefore included in this analysis (Fig. 3).

**Fig. 3:**
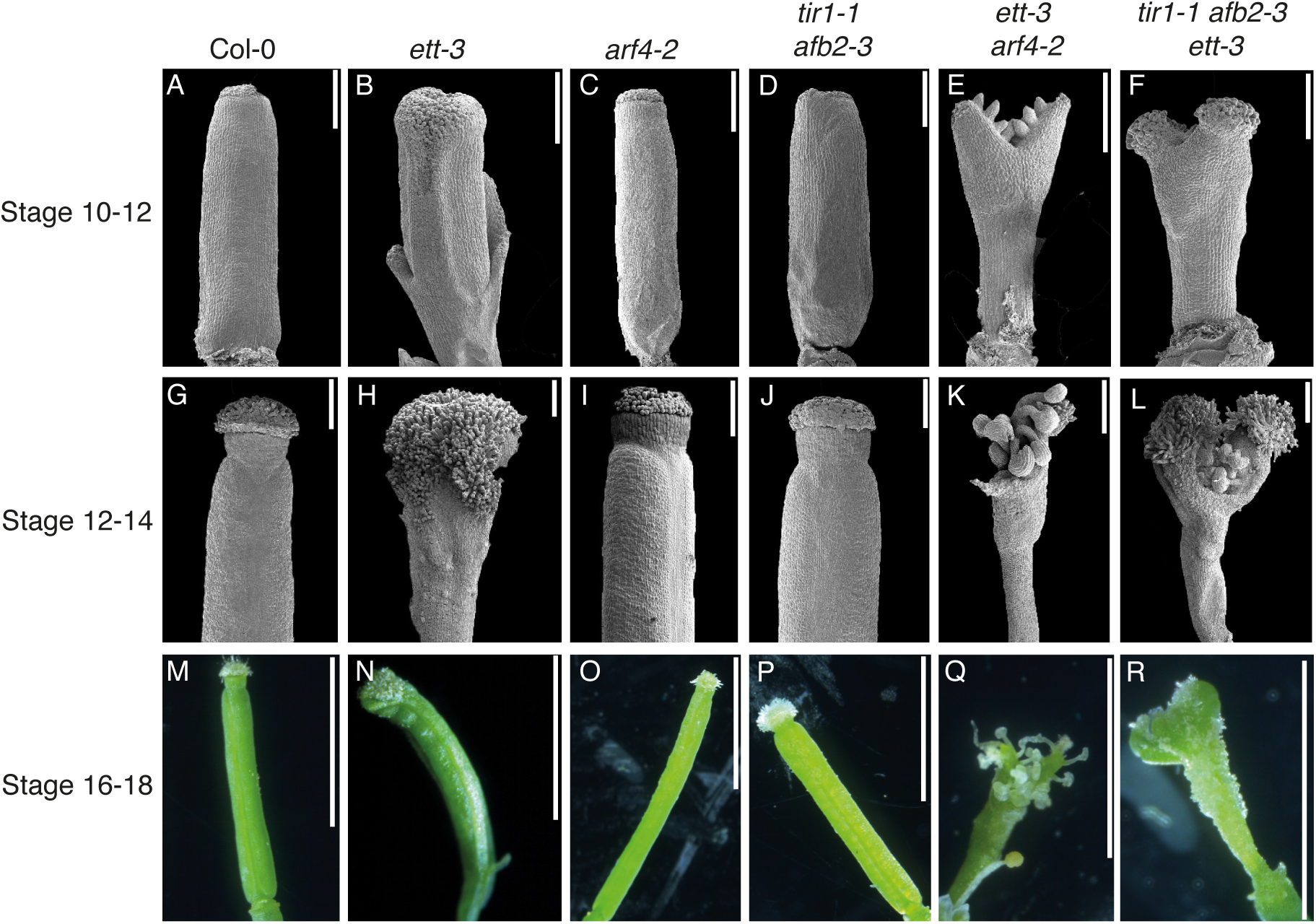
Phenotypic analysis of *Arabidopsis* gynoecium defects in canonical and ETT-mediated pathway mutants reveals synergism in gynoecium development. Scanning electron micrographs (stage 10-12, stage 12-14), and photographs (stage 16-18) of gynoecia from WT (A, G, M), *ett-3* (B, H, N), *arf4-2* (C, I, O), *tir1-1 afb2-3* (D, J, P), *ett-3 arf4-2* (E, K, Q), and *tir1-1 afb2-3 ett-3* (F, L, R) mutants in *Arabidopsis*. Scale bars in SEM 200μm, Scale bars in photographs 3cm

The *ett-3* mutant (Fig. 3B,H,N) exhibited reduced valves, a loss of radial symmetry in the style, and over-proliferation of stigmatic tissues at later developmental stages (Fig. 3H,N), whilst the *arf4-*2 (Fig. 3C,I,O) and *tir1-2 afb2-3* (Fig. 3D,J,P) mutants resembled wild type (Fig. 3A,G,M). The *ett-3* phenotype (Fig. 3B,H,N) was exacerbated in both *ett-3 arf4-2* (Fig. 3E,K,Q) and *tir1-1 afb2-3 ett-3* (Fig. 3F,L,R). Valves in *ett-3 arf4-2* gynoecia were significantly reduced, producing an open ovary with ectopic ovules and few stigmatic papillae (Fig. 3E,K,Q). The *ett-3 arf4-2* gynoecia lacked a style and split into two distinct lobes with stigmatic papillae at their tips (Fig. 3E,K,Q). Similarly, *tir1-1 afb2-3 ett-3* mutants exhibited enhanced valve reduction relative to *ett-3* mutants, producing a partially open ovary with ectopic ovules, and no discernable style or replum (Fig. 3F,L,R). The stigmatic papillae in *tir1-1 afb2-3 ett-3* gynoecia also developed atop two lobes produced at the apex, and exhibited over proliferation of stigmatic tissues at later developmental stages comparable to those observed in *ett-3* (Fig. 3B,F,H,L,N,R).

The exacerbation of the *ett-3* phenotype by *tir1-1 afb2-3* mutation demonstrates that the canonical and ETT-mediated pathways function synergistically to promote gynoecium development. Furthermore, the phenotypic similarities between *ett-3 arf4-2* and *tir1-1 afb2-3 ett-3* mutants, especially at the early stages of gynoecium development, suggest that ETT/ARF4 and TIR1/AFB2 regulate similar processes in gynoecium development, and further implicate ARF4 in the canonical pathway.

To assess whether synergism between the canonical and ETT-mediated pathways is conserved in other *Brassicaceae* species, the same mutant combinations were generated by CRISPR/Cas9 in *Capsella* (Fig. 4). As in *Arabidopsis*, the *Capsella arf4* mutants (Fig. 4C,I,O) resembled wild type (Fig. 4A,G,M), whilst *Crett* displayed valve reduction, stigmatic over proliferation and breakage in the radial symmetry of the style (Fig. 4B,H,N). Whereas the *Arabidopsis tir1-1 afb2-3* mutant phenocopied wild type, valve reduction was observed in the *Crtir1afb2* mutant, and this persisted at later developmental stages so that the shoulders of the heart never fully elongated (Fig. 4,D,J,P). This suggests a more prominent role for the canonical pathway in valve outgrowth in *Capsella* than in *Arabidopsis* and implicates the canonical pathway in heart-shaped fruit development.

**Fig. 4:**
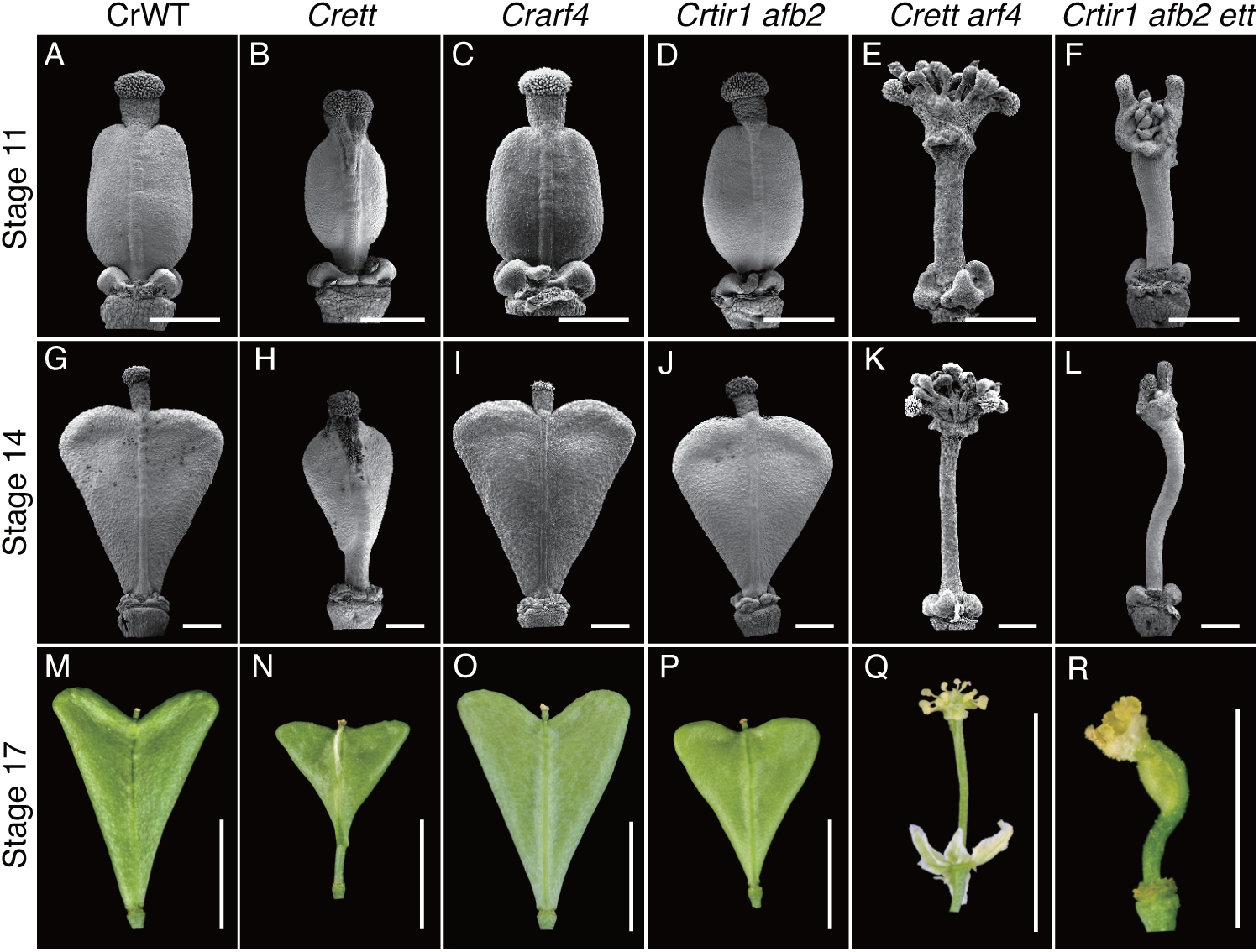
Phenotypic analysis of *Capsella* gynoecium defects in canonical and ETT-mediated pathway mutants reveals conserved synergism in gynoecium development. Scanning electron micrographs (stage 11, stage 14), and photographs (stage 17) of gynoecia from WT (A, G, M), *ett* (B, H, N), *arf4* (C, I, O), *tir1 afb2* (D, J, P), *ett arf4* (E, K, Q), and *tir1 afb2 ett-* (F, L, R) mutants in *Capsella,* Scale bars in SEM 200 μm, Scale bars in photographs 0.5 cm

In *Arabidopsis*, the *ett-3 arf4-2* mutant exhibited the strongest phenotypes, producing a completely open ovary, whilst *the tir1-1 afb2-3 ett-3* mutants exhibited less severe valve reduction defects (Fig. 3). However, *Crettarf4* (Fig. 4E,K,Q) and *Crtir1afb2ett* (Fig. 4F,L,R) mutants exhibited phenotypes which were similar in severity. Both lines produced open ovaries with ectopic ovules atop a hyperextended stalk like structures (Fig. 4E,F,K,L,Q,R). As in *Arabidopsis*, stigmatic tissues formed at the apex of structures at the lateral edges of the gynoecium (Fig. 4E,F,K,L). Taken together these data suggest that the synergism between canonical and ETT-mediated pathways in gynoecium development is conserved between *Arabidopsis* and *Capsella* and may therefore be broadly relevant for gynoecium development across the *Brassicaceae* family.

### Comparative transcriptomics in *Arabidopsis* dissect distinct and overlapping targets of ETT and ARF4 in gynoecium development

ETT is known to function partially redundantly with ARF4 in the regulation of gynoecium development (Pekker et al., 2005; Finet et al., 2010; Kelley et al., 2012). Whilst ETT-mediated signalling and the regulation of ETT-target genes are relatively well studied in the gynoecium (Simonini et al., 2016; Simonini et al., 2017; Kuhn et al., 2020), a global understanding of how ARF4 regulates its targets, and which genes it modulates in conjunction with ETT, is lacking. The phenotypic similarities between *ett arf4* and *tir1 afb2 ett* mutant gynoecia, along with the observation that ARF4 physically interacts with AUX/IAAs (Vernoux et al., 2011), implicate ARF4 in the canonical pathway.

To better understand how ARF4 regulates its targets both alone and together with ETT, a comparative transcriptomic analysis (RNA sequencing, from here on RNA-Seq) was conducted utilizing whole inflorescence tissues of *Arabidopsis* Col-0, *ett-3, arf4-2, and ett-3 arf4-2* lines treated with 100 μM Indole-3-Acetic Acid (IAA) and 10 μM N-1-Naphthylphthalamic Acid (NPA) or mock treatments for one hour.

A principal component analysis revealed that the lines differed more by genotype than by auxin treatment (Fig. S2), and many genes which are reportedly auxin responsive (Paponov et al., 2008) were modulated in response to auxin in wild type plants, indicating that the auxin treatment was effective (Table S1). *PINOID*, a previously well-characterized ETT target (Simonini et al., 2016; Simonini et al., 2017; Kuhn et al., 2019) which is upregulated in wild type in response to auxin (Kuhn et al., 2020) was not identified as an auxin sensitive gene within this study. Previous transcriptomics analyses aiming to identify ETT targets utilized different schedules of auxin treatment (Kuhn et al., 2020; Simonini et al., 2017) which may explain the observed discrepancies. Significant overlap was however observed between genes mis-regulated in *ett-3* mutants between this study and Simonini et al., 2017 (Fig. S3; Table S2 and S3). The data were analyzed to identify auxin independent ETT/ARF4 targets in accordance with the schema in Fig. S4. Two datasets each consisting of three lists of genes were used for this analysis. The first dataset comprised lists of genes which were differentially expressed relative to wild type in *ett-3*, *arf4-2,* and *ett-3 arf4-2* after mock treatment, whilst the second dataset comprised the same after IAA treatment. These lists were compared to generate two three-way Venn diagrams, one for each dataset. The data were then compared across mock and auxin treatments to identify genes which are mis-regulated in the same manner in both the presence and absence of auxin. Any auxin sensitive genes within these lists were removed, leaving genes which are regulated by ETT and ARF4 independently of auxin. Based on where these genes positioned within a three-way Venn diagram, models of gene behavior were devised which may explain the observed patterns of statistical significance (Fig. 5; Fig. S5).

**Fig. 5:**
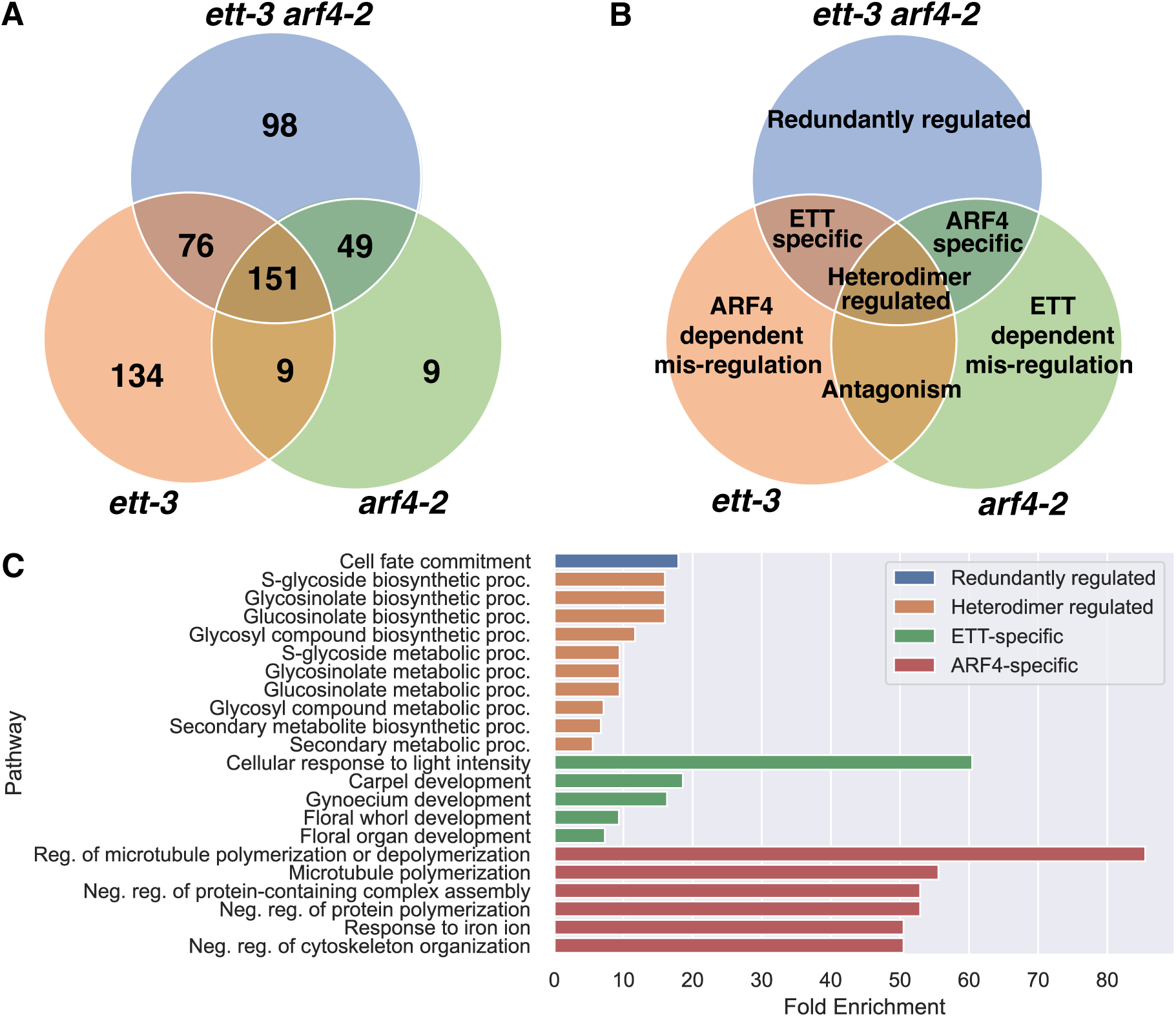
Summary of genes identified in comparative-RNA seq experiments as ETT/ARF4 targets regulated independently of auxin. A) The numbers of genes falling within each segment of a three way venn diagram comparing genes which are significantly differentially independently of auxin in ett-3, arf4-2 and ett-3 arf4-2 mutants compared to wildtype. B) Summary of the relationship between ETT/ARF4 and the genes within each Venn diagram segment. C) Significant GO terms identified for lists of genes in each Venn diagram segment.

Gene Ontology (GO) term analyses were conducted in Shiny GO (Version 0.77, Ge et al., 2020) to identify biological processes which are enriched within each target category (Fig. 5) compared to a background list composed of all genes expressed in the RNA-seq experiment (Supplemental data File 2).

### ETT and ARF4 redundantly regulate cell fate commitment independently of auxin in *Arabidopsis*

The transcriptomics analysis identified 98 genes (53 up, 45 down) which were significantly mis-regulated in *ett-3 arf4-2* relative to wild type in accordance with their redundant regulation by ETT and ARF4 independently of auxin (Fig. 5A,B; Fig S5). GO term analysis indicated significant enrichment within this list for genes involved in cell fate commitment, (Table 1; Fig. 5), including *CLAVATA3/ESR-RELATED 19* (*CLE19*), *NO TRANSMITTING TRACT* (*NTT*), and *FAMA*, which were significantly upregulated; and *YABBY2* (*YAB2*) and *YAB5,* and *ATMYC1* which were significantly downregulated in *ett-3 arf4-2* (Fig. 5C; Fig. 6; Table 1).

**Fig. 6:**
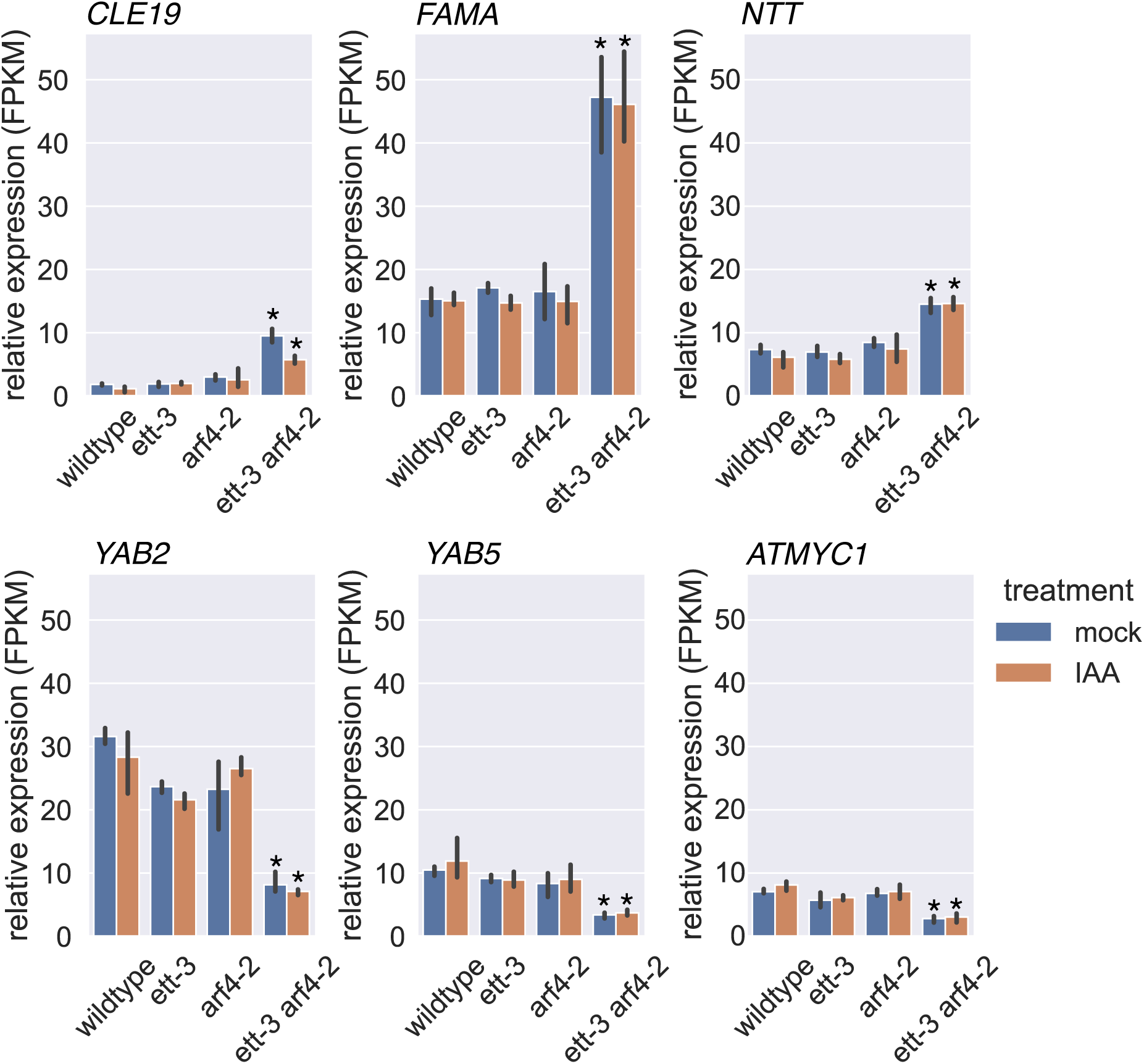
The normalized expression of genes which are redundantly regulated by ETT and ARF4 independently of auxin, and which were identified in GO term analysis as genes involved in the regulation of cell fate commitment. *CLE19*, *CLAVATA3/ESR-RELATED 19, NTT, NO TRANSMITTING TRACT, YAB2, YABBY2, YAB5, YABBY5, ATMYC1, ARABIDOPSIS THALIANA MYC1.* FPMK, fragments per kilobase per million mapped fragments. Statistical analysis was conducted using read count normalization, negative binomial distribution analysis and FDR correction using the BH procedure. The error bars show +/- standard deviation * = Padjust<0.05 vs the same treatment in wildtype.

**Table 1:** Genes which are significantly differentially expressed in *ett-3 arf4-2* mutants relative to wild type both in the presence and absence of auxin, consistent with their redundant regulation by ETT and ARF4, and whose mis-regulation correlates with phenotypes observed in this line.

| Locus | Name | Log2Fold vs WT mock in *ett-3 arf4-2* mock | Log2Fold vs WT IAA in *ett-3 arf4-2* IAA | Significant in all treatments | GO term |
| --- | --- | --- | --- | --- | --- |
| AT3G24225 | *CLE19* | 2.39 | 2.27 | True | cell fate commitment |
| AT3G57670 | *NTT* | 1.01 | 1.22 | True | cell fate commitment |
| AT2G26580 | *YAB5* | -1.6 | -1.7 | True | cell fate commitment |
| AT3G24140 | *FAMA* | 1.66 | 1.57 | True | cell fate commitment |
| AT1G08465 | *YAB2* | -1.94 | -2.04 | True | cell fate commitment |
| AT4G00480 | *ATMYC1* | -1.29 | -1.46 | True | cell fate commitment |
| AT1G70560 | *TAA1* | 1.61 | 1.34 | True | Indole-acetic acid biosynthetic process |
| AT5G03790 | *ATHB-51* | 1.15 | 1.13 | True | floral<br>meristem<br>determinacy |
| AT2G33880 | *WOX9* | 1.66 | 1.90 | True | regulation of<br>meristem<br>growth |
| AT1G80080 | *TMM* | -1.04 | -1.09 | True | stomatal<br>complex<br>formation |

The YABBY transcription factors, *YAB2* and *YAB5*, which are implicated in abaxial cell fate specification, were constitutively downregulated in the double mutant (Table 1; Fig. 6), and this correlated with the abaxial defects overserved in the leaves (Fig. S6 A-C), and patterning defects observed in the gynoecium of this line (Fig. 3).

*FAMA,* a regulator of guard cell differentiation in the stomatal lineage, was upregulated in *ett-3 arf4-2*, and *TOO MANY MOUTHS* (*TMM*), another stomatal regulator, was constitutively downregulated (Table 1). Along with the mis-regulation of stomatal lineage genes in *ett-3 arf4-2*, stomatal defects were also observed on the abaxial surface of *ett-3 arf4-2* leaves (Fig S6). Finger-like protrusions, which resemble the midvein in wild type plants, were produced from the abaxial surface of *ett-3 arf4-2* leaves (Fig. SB,C). In wild type, the midvein lacks stomata, but the projections in *ett-3 arf4-2* exhibited stomata across the surface of these structures (Fig. S6C).

The WUSCHEL related homeobox transcription factor *WOX9*, the CLAVATA3-related peptide encoding gene *CLE19,* and the gene encoding HD-ZIP I transcription factor *ARABIDOPSIS THALIANA HOMEOBOX 51* (*ATHB-51*), were constitutively upregulated in *ett-3 arf4-2* (Table 1); these genes are implicated in meristematic regulation. The mis-regulation of these genes correlates with variability in the numbers of floral organs produced within *ett-3 arf4-2* flowers (Fig. S6D-H), suggesting a role for ETT and ARF4 in floral meristem regulation.

The auxin biosynthesis gene *TAA1*, which promotes the production of auxin through the IPA pathway, was constitutively upregulated in the double mutant (Table 1), suggesting that auxin biosynthesis is also regulated redundantly by ETT and ARF4 independently of auxin.

Together, these data correlate with phenotypes observed in the *ett-3 arf4-2* mutant and suggest that ETT and ARF4 may redundantly regulate cell fate specification independently of auxin in a range of biological contexts, including axis specification of lateral organs.

### Transcriptomics identify ETT and ARF4-specific targets in *Arabidopsis*

Transcriptomics revealed 76 targets which became similarly mis-regulated in both *ett-3* and *ett-3 arf4-2*, consistent with their ETT-specific regulation (Fig. 5A,B; Fig. S5). The list of ETT-specific targets was enriched in GO term analysis for genes involved in gynoecium development, carpel development, floral whorl development, floral organ development, and cellular response to light intensity (Table 2; Fig. 5C; Fig. 7). The list included zinc finger transcription factor *JAGGED* (*JAG*), the YABBY transcription factor *CRABSCLAW* (*CRC*), and basic-Helix-Loop-Helix TF *INDEHISCENT (IND)* which have previously been implicated in gynoecium development (Table 2; Fig. 7). *JAG* expression was constitutively downregulated in *ett-3* and *ett-3 arf4-2* mutants relative to wild type independently of auxin. JAG functions with NUBBIN (NUB) to regulate gynoecium development (Dinneny et al., 2006). As in *ett-3* and *ett-3 arf4-2*, *jag nub* mutants show apical defects in the gynoecium and altered floral organ development (Fig. 3; Fig. S6D-H). *CRC* was constitutively downregulated in *ett-3* and *ett-3 arf4-2*, and the knockdown of this gene has previously been shown to cause apical gynoecium defects in *Arabidopsis* (Bowman & Smyth, 1999). The mis-regulation of these genes therefore correlates with phenotypes observed in the *ett-3* and *ett-3 arf4-2* mutant gynoecia.

**Fig. 7:**
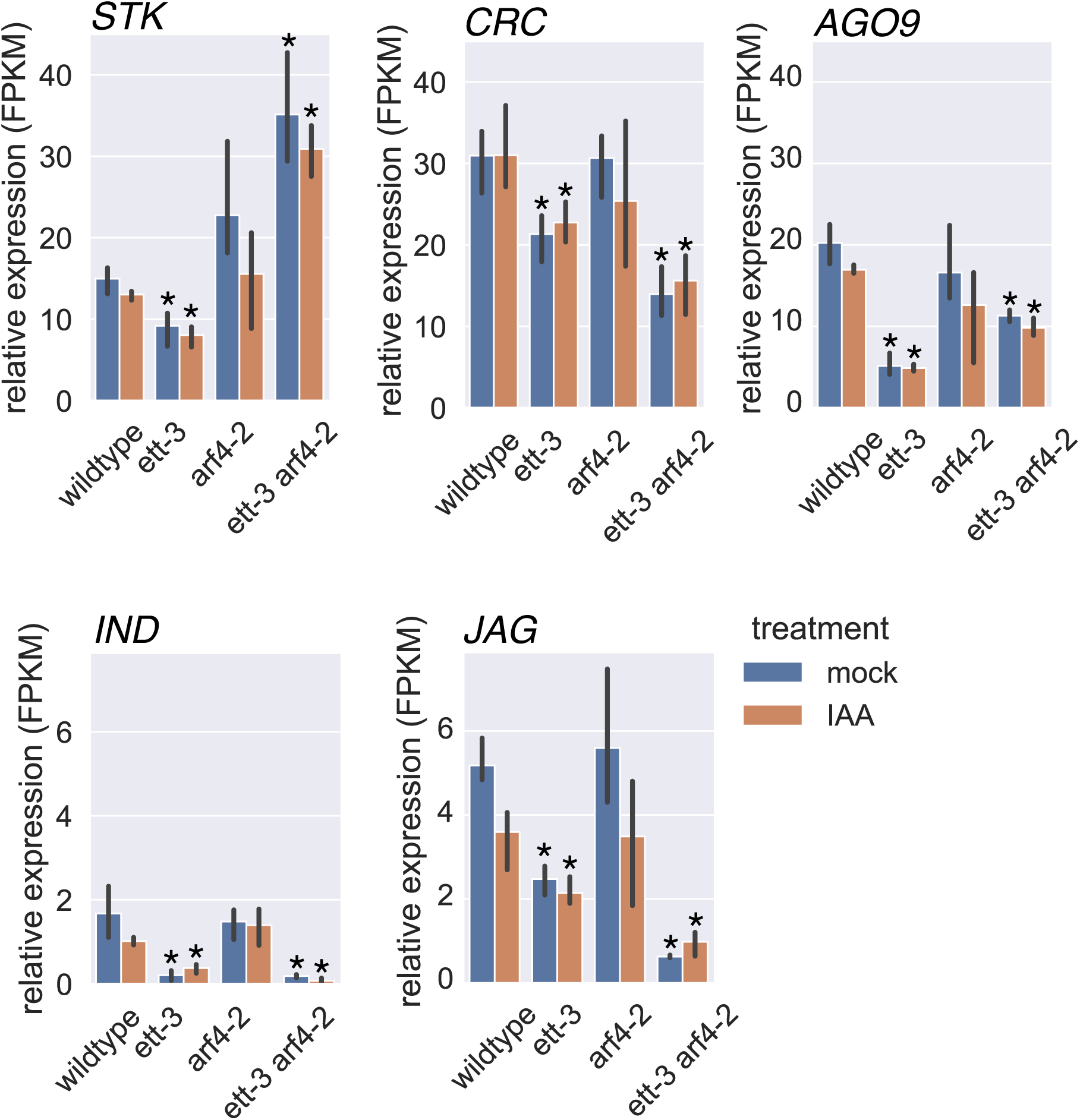
The normalized expression of genes identified in GO term analysis as involved in gynoecium development which are differentially expressed in *ett-3* and *ett-3 arf4-2* relative to wildtype, consistent with their ETT-specific regulation. *STK*, *SEEDSTICK*, *CRC*, *CRABSCLAW*, *AGO9*, *ARGONAUT9*, *IND*, *INDEHISCENT*, *JAG*, *JAGGED*. FPMK, fragments per kilobase per million mapped fragments. Statistical analysis was conducted using read count normalization, negative binomial distribution analysis and FDR correction using the BH procedure. The error bars show +/- standard deviation * = Padjust<0.05 vs the same treatment in wildtype.

**Table 2:** Genes which are significantly differentially expressed in *ett-3* and *ett-3 arf4-2* mutants relative to wild type both in the presence and absence of auxin, consistent with their ETT-specific regulation.

| Locus | Name | Col-0 vs<br>*ett-3*<br>mock<br>log2fold<br>change | Col-0 vs<br>*ett-3* IAA<br>log2 fold<br>change | Col-0 vs<br>*ett-3 arf4-2*<br>mock<br>log2fold<br>change | Col-0 vs<br>*ett-3 arf4-2*<br>IAA<br>log2fold<br>change | Significance<br>In all groups | Go Terms |
| --- | --- | --- | --- | --- | --- | --- | --- |
| AT1G69180 | *CRC* | -0.44 | -0.42 | -1.12 | -1.02 | True | Carpel<br>Development |
| AT5G21150 | *AGO9* | -1.87 | -1.78 | -0.82 | -0.83 | True | Floral organ<br>development |
| AT4G09960 | *STK* | -0.60 | -0.68 | 1.27 | 1.21 | True | Carpel<br>Development |
| AT3G22840 | *ELIP1* | 1.63 | 1.75 | 1.41 | 1.35 | True | Response to<br>light intensity |
| AT1G29440 | *SAUR63* | -1.18 | -1.43 | -1.08 | -1.15 | True | Floral organ<br>development |
| AT1G68480 | *JAG* | -0.96 | -0.72 | -2.97 | -1.88 | True | Carpel development |
| AT4G14690 | *ELIP2* | 1.54 | 1.13 | 1.56 | 1.14 | True | Response to light intensity |
| AT4G00120 | *IND* | -2.82 | -1.38 | -3.03 | -3.61 | True | N/A valve margin specification (Liljegren et al., 2004) |

Transcriptomics revealed several groups of genes regulated by ETT and ARF4 which did not appear to regulate gynoecium development. This included 134 genes mis-regulated only in *ett-3* single mutants, nine genes mis-regulated only in *arf4-2* single mutants, and nine genes mis-regulated in both *ett-3* and *arf4-2* single mutants (Fig. 5A,B), which were not enriched in GO term analysis, and are not discussed further.

Whilst the *arf4-2* mutant appeared as wild type (Fig. 3), 49 ARF4-specific genes were identified (Fig. 5A,B; Fig. S5). This list was enriched for genes involved in the regulation of microtubule polymerization and related GO terms (Fig. 5C) and did not contain any known regulators of gynoecium development, suggesting that ARF4 mainly functions in conjunction with ETT to elicit its auxin-independent effects in the carpels.

151 genes were significantly mis-regulated in all mutants relative to wild type, consistent with their regulation by ETT/ARF4 heterodimers (Fig. 5A,B; Fig. S5). This list was significantly enriched in GO term analysis for genes involved in secondary metabolism including S-glycoside, glucosinolate and glycosinolate biosynthesis (Fig. 5C).

Together, the transcriptomics data suggest that ETT and ARF4 regulate distinct and overlapping targets independently of auxin to promote gynoecium development and imply that ARF4 mainly functions with ETT to promote its auxin independent effects on gynoecium development.

### Auxin sensitive analysis in *Arabidopsis* suggests redundant maintenance of auxin insensitivity by ETT and ARF4

Gynoecium morphogenesis is dependent on the dynamic distribution of auxin (Moubayidin & Østergaard, 2014), and gynoecium defects of *ett-3 arf4-2* mutants suggest a role for ETT and ARF4 in the transduction of auxin signalling (Pekker et al., 2005). A second analysis was therefore conducted in *Arabidopsis* to assess how the auxin response varies from wild type in the mutants. Genes which were significantly auxin responsive in wild type were compared to genes which were significantly auxin responsive in each mutant background (*ett-3*, *arf4-2,* and *ett-3 arf4-2*).

Few genes were identified in the auxin sensitive analysis with a known role in gynoecium development. 179 genes were auxin sensitive only in the *ett-3 arf4-2* mutant background (44 up and 135 down upon auxin treatment, Fig. 8) consistent with their redundant regulation by ETT and ARF4, which maintain their auxin insensitivity. This list was enriched for a range of GO terms including pectin metabolism/catabolism (Fig. 9A). The mis-regulation of pectin methylesterification has previously been linked to gynoecium defects in *ett* mutants (Andres-Robin et al., 2018), and several pectin methlyesterases (PMEs) and pectin methylesterase inhibitors (PMEIs) have been previously identified as direct ETT targets (Andres-Robin et al., 2020). The list of redundantly regulated ETT/ARF4 targets in the auxin sensitive analysis conducted here included *PME23*, *PME28*, *PME48*, *PME49, and* members of the pectin lyase superfamily (*AT1G05660, AT3G07850, AT5G48140, AT3G07830*, *AT3G07840, AT3G14040*).

**Fig. 8:**
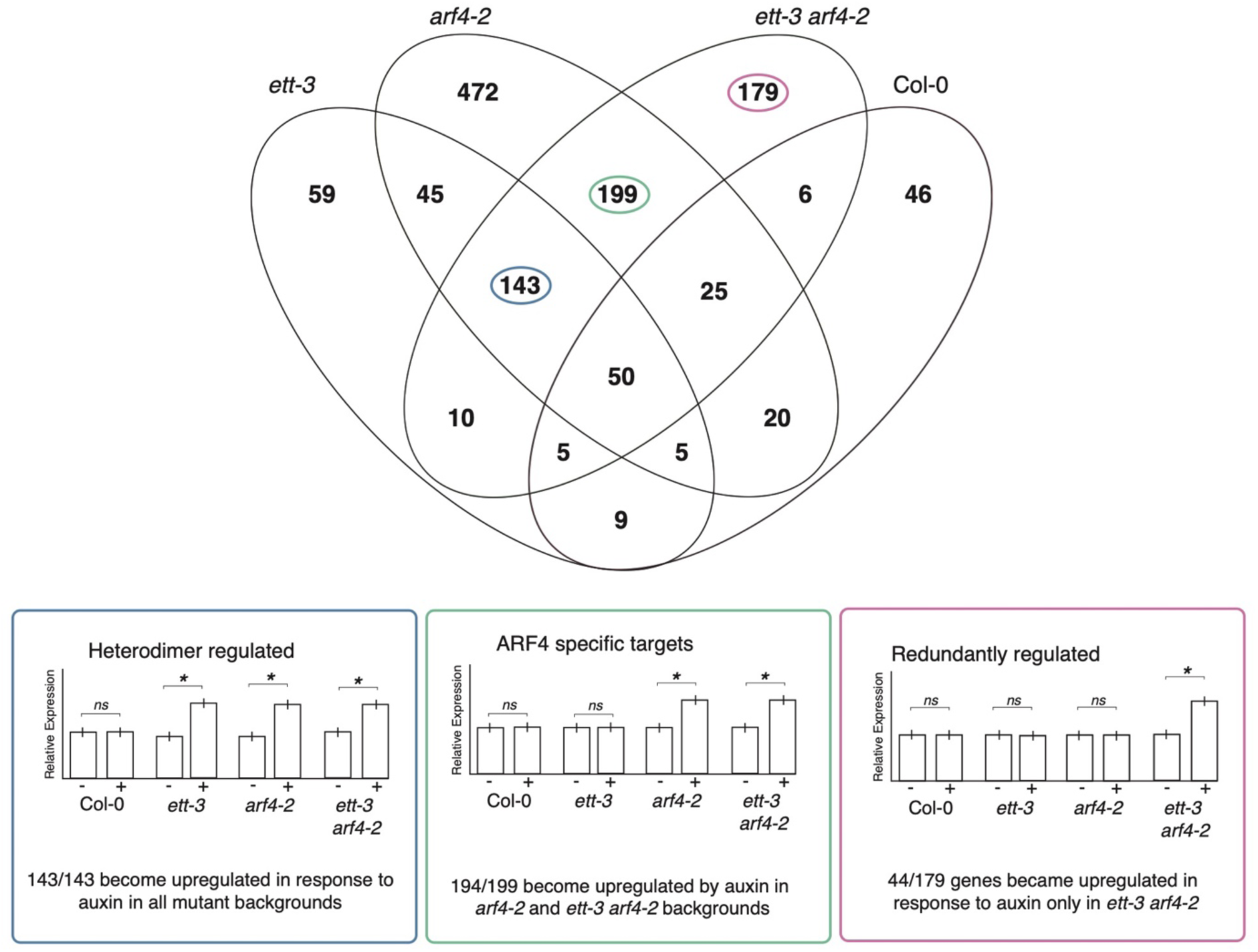
Summary of the outcome of the auxin sensitive comparative RNA-sequencing analysis comparing the auxin response in ett-3, arf4-2, or ett-3 arf4-2 to the auxin response in wildtype. Groups of genes which conform to models of A-B ARF antagonism, and their expression profiles are highlighted below.

**Fig. 9:**
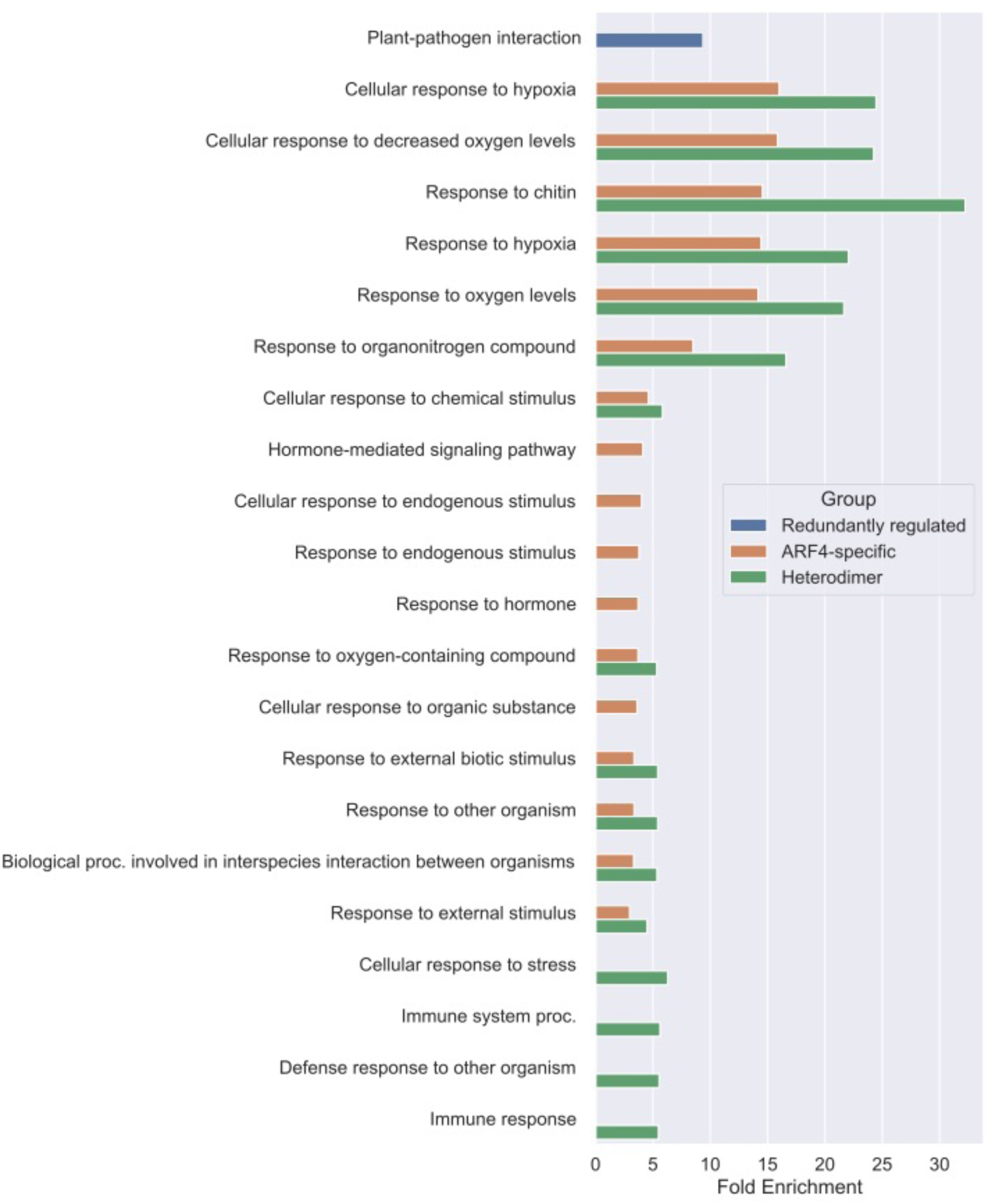
GO terms which are significantly enriched in lists of genes which conform to models of A-B ARF antagonism.

None of the other gene lists assessed in the auxin sensitive analysis contained known regulators of gynoecium development, but the data did implicate ETT and ARF4 in the maintenance of auxin insensitivity at target loci involved in a range of biological processes, which then become auxin sensitive upon ETT/ARF4 mutation consistent with previously proposed models of A-B ARF antagonism.

### Auxin sensitive analysis reveals patterns of gene regulation consistent with A-B ARF antagonism

Previous studies in *Marchantia polymorpha* and *Phiscomitrium patens* found antagonism between A- and B-ARFs to regulate the auxin sensitivity of tissues (Kato et al., 2020; Lavy et al., 2016), with the auxin response being promoted by A-ARFs and inhibited by B-ARFs. Within the auxin sensitive analysis several groups of genes were identified which behave in a manner consistent with A-B ARF antagonism (Fig. 8). Firstly, 44 genes (out of 179) became upregulated in response to auxin only in *ett-3 arf4-2* mutants, suggesting that ETT and ARF4 act redundantly to prevent their auxin responsiveness (Fig. 8). Secondly, a further 143 genes (out of 143) were auxin insensitive in wild type and upregulated in response to auxin in all mutant backgrounds, suggesting that ETT-ARF4 dimers maintain auxin insensitivity at these loci (Fig. 8). Interestingly, ETT and ARF4 showed physical interactions in Yeast 2-Hybrid (Y2H) assays (Fig. S7). ETT-ARF4 interaction was dependent on the presence of the ARF4 DNA binding domain and was not sensitive to auxin (Fig. S7). Together the Y2H and RNA seq data suggest that ETT and ARF4 can form heterodimers to regulate gene expression.

Finally, 194 genes (out of 199) were auxin insensitive in wild type and became upregulated upon auxin treatment in *arf4-2* and *ett-3 arf4-2* mutants, consistent with the ARF4-specific prevention of their auxin sensitivity (Fig. 8). GO term analysis showed that list of 44 redundantly regulated genes which become auxin responsive in *ett-3 arf4-2* mutants are involved in plant-pathogen interactions. The list of heterodimer targets which became auxin responsive in all mutant backgrounds were enriched for loci involved in responses to environmental stresses, including biotic stimuli and hypoxia (Fig. 9). The list of ARF4-specific targets was also enriched for similar GO terms (Fig. 9). Together these data suggest a role for ETT and ARF4 in the regulation of a range of responses to biotic and abiotic stress. Consistent with the hypothesis that ETT and ARF4 may antagonise A-ARFs, *ARF5*, *ARF6,* and *ARF8*, which are A-class activators, are expressed in the gynoecium at various stages of development (Fig.S8), however future studies will be required to assess whether they share antagonistic roles to ETT and ARF4 in the regulation of environmental stress.

## Discussion

### Synergism between canonical and ETT-mediated signalling components in gynoecium development

The phenotypic and expression analyses presented here suggest the existence of synergistic regulation between the canonical and ETT-mediated pathways in gynoecium development, and that this mechanism is conserved between *Arabidopsis* and *Capsella*. Indeed, previous studies in both *Arabidopsis* and *Capsella* have shown that the dynamic distribution of auxin plays an important role in gynoecium development (Moubayidin and Østergaard, 2014; Dong et al., 2019).

Whilst a role for ETT in style development has been observed in *Brassica rapa* (Simonini et al., 2018), ETT’s role in *Capsella* fruit development had not been characterized prior to this study. In *Arabidopsis* ETT was known to function partially redundantly with ARF4 in gynoecium development (Pekker et al., 2005; Finet et al., 2010). Here, we show that as in *Arabidopsis*, ETT and ARF4 function partially redundantly in *Capsella*; phenotypes present in *Crett* mutants are exacerbated in *Crett arf4*, suggesting that ARF4 shares some, but not all, of its functionality with ETT.

Furthermore, the *tir1 afb2 ett* triple mutant phenocopies *ett arf4* mutants in both species studied, suggesting a role for ARF4 in the integration of signalling with the canonical pathway. Consistent with this hypothesis, ARF4 has previously been shown to interact with AUX/IAAs of the canonical pathway through its C-terminal domain (Vernoux et al., 2011). Structural evolution studies have previously indicated that ETT and ARF4 arose from the duplication of a non-truncated ARF prior to the radiation of extant angiosperms (Finet et al., 2010), and that the truncation of either ETT or ARF4, so that they lack the C-terminal PB1 domain, is common in the angiosperm lineage. However, whilst *Arabidopsis* ETT was more able to rescue *ett* phenotypes when truncated, ARF4 required its C-terminal domain to function (Finet et al., 2010), consistent with the idea that AUX/IAA interaction is required for ARF4 activity. However, the C-terminal PB1 domain has also been shown to facilitate ARF homodimer and homo-oligomerization (Davis et al., 2020; Korasick et al., 2015), meaning that further studies will be required to understand the mechanism of ARF4 action, and how it integrates the canonical and ETT-mediated pathways.

### Comparative transcriptomics correlate with phenotypic defects observed in *Arabidopsis* mutants

The auxin sensitive analysis identified distinct and overlapping targets of ETT and ARF4. Targets under the auxin-independent redundant regulation of ETT and ARF4 included the YABBY TFs *YAB2* and *YAB5*, whilst *CRC* was mis-regulated in both *ett-3* and *ett-3 arf4*-2, consistent with its ETT-specific regulation. YABBY proteins specify abaxial identity along with the KANADI family of transcription factors (Eshed et al., 2004). ETT and ARF4 function co-operatively with KANADI proteins to promote abaxial cell fate of lateral organs (Pekker et al., 2005). Laminal outgrowths from the abaxial leaf surface have previously been observed in *kan1 kan2* loss of function mutants. This phenotype is suppressed in *yab1 yab3 yab5* (Eshed et al., 2004) and *yuc1 yuc2 yuc4* backgrounds (Wang et al., 2011), suggesting that the production of outgrowths is dependent on YABBYs and auxin biosynthesis. Similar outgrowths were observed on the abaxial surface of *ett arf4 arf2* triple mutants (Guan et al., 2017) and the *ett-3 arf4-2* mutant leaves in this study, consistent with previously proposed roles for ETT/ARF4 in KANADI-mediated signalling (Pekker et al., 2005). However, given that *YAB2*, *YAB5,* and *CRC* were significantly downregulated in the *ett-3 arf4-2* mutant background, it is unclear if these outgrowths are YABBY dependent. Whilst *yabby* single mutants lack obvious abaxial-adaxial identity defects, higher order mutants show significant alterations in abaxial-adaxial polarity (Sarojam et al., 2010; Siegfried et al., 1999). The downregulation of multiple *YAB* transcription factors within the *ett-3 arf4-2* mutant is therefore consistent with the abaxial-adaxial identity defects observed in the leaves and gynoecia of the double mutant in this study.

Previous studies reported reduced valve margin definition in *ind* (Liljegren et al., 2004), and apical gynoecium defects in *jag nub* and *crc* mutants (Dinneny et al., 2006; Bowman and Smyth, 1999). Similar gynoecium defects were exhibited in this study by *ett-3* and *ett-3 arf4-2* (Fig. 1), along with the constitutive downregulation of *IND, JAG,* and *CRC,* which is consistent with the previously proposed roles of these genes in gynoecium development. Future studies should aim to determine which of the targets identified within this study are direct/indirect ETT/ARF4 targets.

### Auxin Sensitive analysis suggests ETT and ARF4 maintain auxin insensitivity at target loci

Previous studies in *Marchantia polymorpha* and *Phiscomitrium patens* found antagonism between A- and B-ARFs to regulate the auxin sensitivity of tissues (Kato et al., 2020; Lavy et al., 2016), with the auxin response being promoted by A-ARFs and inhibited by B-ARFs. Consistent with this model of B-ARF function, ETT and ARF4 were previously shown to repress auxin responses in an *Arabidopsis* root hair system (Choi et al., 2018). The transcriptomics data presented here suggest that ETT and ARF4 largely prevent the auxin sensitivity of distinct and overlapping targets, many of which become upregulated in response to auxin in various mutant backgrounds, consistent with models of A/B-ARF antagonism.

However, many of their redundantly regulated targets, including genes involved in the regulation of pectin become downregulated in response to auxin. Auxin-mediated repression has been observed in TMK-mediated signalling, where atypical AUX/IAAs are stabilized in the presence of auxin (Cao et al., 2019). However, it is unclear if such a mechanism functions in the gynoecium. Previous studies have implicated PME family members in the regulation of cell wall stiffness during gynoecium development in *ett* mutants and showed that ETT promotes PME expression in the carpels (Andres-Robin et al., 2018; Andres-Robin et al., 2020). Furthermore, the overexpression of *PECTIN METHYLESTERASE INHIBITOR 3* (*PMEI3*), which antagonises PME activity, phenocopied the valve reductions observed in *ett* mutants (Andres-Robin et al., 2018). Taken together with these previous studies, the data presented here suggest that ETT and ARF4 may redundantly promote the expression of *PMEs* to prevent their downregulation in response to auxin, and the mis-regulation of these genes may contribute to the gynoecium defects observed in *ett-3 arf4-2*.

The data presented suggest that ETT functions alongside ARF4 to prevent the expression of genes which would otherwise become auxin sensitive in its absence, possibly through the activity of A-ARFs with which they are co-expressed, or due to the activity of TMK-related pathways in the case of auxin-downregulated genes. The data also suggest that the maintenance of genes in an auxin insensitive state may be equally as important as the promotion of auxin sensitivity at other loci to facilitate normal growth and development. This study demonstrates how dissecting the relationships between distinct auxin signalling pathways is a powerful approach to gain a greater understanding of the wider auxin signalling network, thereby broadening the perspective of auxin-mediated plant development.

## Materials and Methods

### Plant materials and growth condition

#### Arabidopsis

Seeds were sown onto damp soil (Levington F2 compost with Intercept and grit at a 6:1 ratio) and stratified for 3 days at 4°C in the dark before transferal to a controlled environment room (CER) and growth in long day conditions (16 hours light/8 hours dark, 22°C). All mutant lines were generated in a Col-0 background. Mutant and transgenic lines used in this work which have been previously published are: *ett-3* (Sessions et al., 1997; Simonini et al., 2016), *arf4-2* (SALK_070506, Alonso et al., 2003) and *tir1-1 afb2-*3 (Parry et al., 2009). The *ett-3 arf4-2* line was generated by crossing the *ett-3* line and *arf4-2* (SALK_070506). The *tir1-1 afb2-3 ett-3* line was generated by crossing *ett-3* with *tir1-1 afb2-3*. The PCR conditions and genotyping primers used for these lines are listed in Table S4 and Table S5 respectively. *pARF3/ETT: GFP*, *pARF4: GFP*, *pARF5: GFP*, *pARF6: GFP* and *pARF8: GFP* are previously published (Rademacher et al., 2011). The *pETT:GUS* line was previously published in Kuhn et al., 2020.

#### Capsella

The genetic and expression analysis in *Capsella rubella* was conducted in the Cr22.5 background. The seeds were germinated on MS medium containing 10 µM Gibberellin at 22°C. 10-day-old seedlings were then transplanted to soil in 8cm pots and moved into a controlled environment room (CER) at 22°C, 16 hrs light and 20°C 8 hrs dark conditions.

### Genotyping

Genomic DNA was extracted from fresh young leaves by grinding with a micro-pestle in a 1.7 ml tube. Arabidopsis extraction buffer was added (200 mM Tris HCl pH 7.7, 250 mM NaCl, 25 mM EDTA and 0.5% SDS) and the sample was ground for an additional 30 seconds. The sample was then centrifuged at 15,000 x g in a table-top centrifuge for 5 minutes. The supernatant was transferred to a fresh 1.5 ml tube containing 200 μl isopropanol without disturbing the pellet. The sample was then inverted at room temperature for 2 minutes. The sample then underwent centrifugation at 15,000 x g in a table-top centrifuge for 7 minutes, and the supernatant was discarded. The pellet was then air dried for 20 minutes at room temperature and resuspended in 50 μl dH2O before being used as a template for PCR or frozen at -20°C. SALK lines (*arf4-2 and afb2-3*) were genotyped in accordance with Alsonso et al., 2003 using the primers and conditions specified in Table S4 and S5. Genotyping PCRs were conducted using GoTaq DNA Polymerase (Promega) in accordance with the manufacturer’s guidelines, and with a final volume of 20 μl where genotype assessment occurred through gel electrophoresis (*arf4-2 and afb2-3*), or 50 μl if genotype was to be assessed by sequencing (*ett-3*) or BsaI restriction digestion (*tir1-1*). The *ett-3* EMS mutation causes a G to A substitution at position 2226 of the *AT2G33860* genomic sequence. PCR was conducted using the primers specified in Table S4 and S5, to amplify a 246bp product which was purified with a QIAquick PCR purification kit (Quiagen) before sequencing. The *tir1-1* allele is an EMS mutation which introduces a G to A mutation in position 440 of the genomic *AT3G62980* sequence. A 517bp product was amplified using the primers specified in Table S4 and S5 and digested with BsaI-HFv2 (NEB) according to the manufacturer’s guidelines before gel electrophoresis of the resulting product on 3% agarose gel. BsaI digestion of the PCR product in the *tir1-1* mutant yields a single band of 517bp, whilst the wild type produces two bands of 437 and 80bp.

### Plasmid construction and plant transformation

For the construction of the promoter: GUS reporter plasmids, the promoter regulatory sequences of *CrAFB2* (1941bp), *CrAFB3* (2072bp)*, CrTIR1* (2077bp)*, CrARF4* (5324bp) and *CrETT* (3029bp), *AtAFB2* (1086bp), *AtAFB2* (2049bp) and *AtTIR1* (2076bp) were isolated by PCR from genomic DNA and inserted upstream of *GUS* gene of pCambia1301 vectors. For the construction of the CRISPR/Cas9 mediated genome editing plasmids, DNA sequences encoding gRNAs adjacent to the PAM sequences (NGG) were designed to target the first or second exons of gene of interest. The gRNAs were synthesised as oligonucleotides with golden-gate cloning adapters and then were insert downstream of U6 promoters. The resulting gRNA plasmid were then recombined with *pRPS5a:Cas9z:E9t* and hygromycin selection marker using golden-gate cloning methods to produce the binary vectors. All vectors were verified by sequencing and introduced into *Agrobacterium tumefaciens* strain LBA4404 by electroporation.

Transformation of *Capsella* plants followed the methods described in Dong et al., 2022. The transformants were screened on MS plants with 1% sucrose and 0.8% agar containing 40mg/L hygromycin (Roche, no. 10843555001). For each construct, at least 10 independent transformants were obtained for further analysis. For the CRISPR transformants, preliminary genotyping was conducted using primers listed in the Table S6. Only the plants showing evidence of gene editing were kept and processed into the T2 generation. For generating the Cas9-free homozygous mutant, the T2 CRISPR transformants were screened on MS plants with 1% sucrose and 0.8% agar containing 10mg/L hygromycin. the hygromycin sensitive plants were isolated and genotyped again using primer listed in Table S6. All the analysis were conducted on the T2 generation of the representative transgenic lines. Additional information on the mutations induced in each line is available in Fig.S9.

### Auxin treatment, expression analysis and GUS staining

For the auxin treatment, 100 μM IAA and 10 μM NPA working solutions were prepared with ethanol and tween 20 (5%). The auxin and mock (water with 5% tween 20) solutions were dipped onto the inflorescence after flowers from stage 13 onwards had been removed. The plants were then kept under a vacuum for 5 mins and kept in humid condition in the CER under long-day condition mentioned above for 1h.

The GUS histochemical assays were performed as previously described in Dong et al., 2019. Specifically, inflorescence samples of *Capsella* or *Arabidopsis* were fixed in acetone for 20 mins at - 80°C, and then washed twice for 5 mins in 100 mM sodium phosphate buffer. The samples were then processed in 100 mM sodium phosphate buffer containing 1mM K_3_Fe(CN)_6_, 1mM K_4_Fe(CN)_6_ at room temperature for 30 mins to block the diffusion of GUS proteins. The staining process was conducted at 37°C in the X-Gluc solution for 6-8h. The X-Gluc solution contains 100 mM sodium phosphate buffer, 10mM EDTA, 0.5 mM K_3_Fe(CN)_6_, 3 mM K_4_Fe(CN)_6_, 0.1% Triton X100 and 1 mg/ml of ß-glucoronidase substrate X-gluc (5-bromo-4-chloro-3-indolyl glucuronide, MELFORD: B40020) dissolved in DMSO. After staining, the reaction was suspended and replaced with 70% ethanol several times until chlorophyll was completely washed out. Gynoecia or young fruits were dissected, mounted in Chlorohydrate (Sigma number: 15307-500G-R) solution and analyzed using a Zeiss Axio Imager light microscope.

### Scanning Electron Microscopy

The Scanning Electron Microscopy (SEM) experiment followed the protocol in Dong et al., 2019. Specifically, the gynoecia of each genotype were fixed in FAA and dissected. The samples were critically-point dried in CO2 and sputter-coated with gold. The samples were subsequently examined using a Zeiss Supra 55VP field Scanning Electron Microscope with an acceleration voltage of 3.0 kV.

### RNA sequencing Analysis (*Arabidopsis*)

Whole inflorescences were harvested from Col-0, *ett-3*, *arf4-2,* and *ett-3 arf4-2* lines treated with either mock (dH2O, 0.1% tween, 0.1% ethanol) or 100 μM IAA and 10μM NPA (dissolved in ethanol), 0.1% tween for one hour. Three biological replicates were harvested for each genotype one hour after treatment. Each replicate contained pooled material from three plants. The tissue was snap frozen in liquid nitrogen and RNA extraction was conducted using a Qiagen RNeasy Plant Mini Kit with on-column DNase treatment, in accordance with the manufacturer’s guidelines. The RNA quality was assessed by gel electrophoresis, and the RNA concentration was measured using a nanodrop (ThermoFisher), and the RNA integrity was tested using a Bioanalyzer 2100 (Agilent). Novogene (Hong Kong), conducted cDNA library preparation (250-300 bp insert cDNA library), and sequencing (20M pair reads). The mRNA was enriched using oligo-dT beads and fragmented before cDNA synthesis with reverse transcriptase. First strand synthesis was conducted, and second strand synthesis buffer (containing dNTPs, RNase H and E. coli polymerase I, Illumina) was added so that the second strand was produced by nick-translation. The library was then purified and underwent terminal repair, before A-tailing, ligation of sequencing adapters, size selection and PCR enrichment. Library concentration was assessed using a Qubit 2.0 fluorometer (Life Technologies) and the strand sizes were verified using a Bioanalyzer 2100 (Agilent) and qPCR. Raw Illumina sequencing data was then converted to sequence reads using base calling. The error rate (0.03%) and A/T/G/C Content Distribution were tested as a quality control, before data filtering to remove low quality reads, and reads with adaptor contamination. Sequencing reads were mapped to the reference genome (TAIR10), using HISAT2. Gene expression level was assessed using transcript abundance, which assesses the number of Fragments Per Kilobase of transcript sequence per Millions base pairs sequenced (FPKM). This method accounts for both the effects of sequencing depth and gene length on fragment counting. HTSeq software was utilized in this analysis. Differentially Expressed Gene (DEG) analyses utilized DESeq2 to conduct readcount normalization and negative binomial distribution analysis (p<0.05) followed by the determination of the false discovery rate using the Benjamini and Hochberg procedure (p_adjust_<0.05). The FPKMs of all genes expressed in this analysis are available in Supplemental data File 1.

### Principle Component Analysis

Principle component analysis was conducted in iDEP 9.1 (http://ge-lab.org/idep/, (Ge et al., 2018) on expression values normalised to fragments per kilobase of transcript per million mapped reads (FPKM).

### Go Term Analysis

Gene Ontology (GO) term analysis was conducted on the lists of genes which fell into each category of the Venn diagrams in both the auxin independent and auxin sensitive analyses to identify whether these lists were enriched for genes involved in specific biological processes. This GO term analysis was conducted using shiny GO (version 0.77, Ge et al.,2020) using the default parameters (FDR cut off =0.05). Gene lists were compared to all gene found to be expressed in the RNA seq analysis (Supplemental data File 2)

### Analysis of auxin independent ETT/ARF4 targets

This analysis was conducted using the differentially expressed gene lists provided by Novogene (Hong Kong). Six lists of differentially expressed genes were used in this analysis: (1) *arf4-2* vs Col-0 mock; (2) *ett-3* vs Col-0 mock; (3) *ett-3 arf4-2* vs Col-0 mock; (4) *arf4-2* vs Col-0 IAA; (5) *ett-3* vs Col-0 IAA; and (6) *ett-3 arf4-2* vs Col-0 IAA. Only genes which were at least 2-fold up or downregulated in at least one background were included in this analysis (log2fold value of >1 or < -1). Analyses were conducted with Venny (Version 2.1; https://bioinfogp.cnb.csic.es/tools/venny/index.html; Oliveros, 2007) to identify genes mis-regulated only in one background, mis-regulated only in two backgrounds, or mis-regulated in all three backgrounds relative to wild type. The mock and auxin treated datasets were independently assessed using Venny, and then compared to identify genes which were similarly differentially expressed in both the presence and absence of auxin. Finally, any genes which were significantly differentially expressed (2-fold up or down-regulated in at least one background) in response to auxin were removed from the lists, leaving only those genes which were regulated independently of auxin. The output of this analysis is available in Supplemental data File 3.

### Analysis of auxin sensitive ETT/ARF4 targets

The auxin sensitive analysis utilized DEGs from the following lists generated by Novogene (Hong Kong): (1) Col-0 IAA vs Col-0 mock; (2) *ett-3* IAA vs *ett-3* mock; (3) *arf4-2* IAA vs *arf4-2* mock; and (4) *ett-3 arf4-2* IAA vs *ett-3 arf4-2* mock. The genes which were differentially expressed in each background in response to auxin were then compared to reveal how the auxin response in each mutant differed from WT. Venny (Version 2.1) was utilized to generate a four-way Venn diagram. The output of this analysis is available in Supplemental data File 4.

### Yeast 2 Hybrid

The full-length coding sequence of ETT and ARF4 and truncated ARF4 fragments were amplified by PCR using primers designed for gateway cloning (Supplemental Table 6). The resulting PCR products were purified using a Qiagen QIAquick PCR purification kit and introduced in the gateway entry vector pDONR207 (Life Technologies) by BP clonase reactions, in accordance with the Thermo-Fisher Gateway protocol. The products of this reaction were transformed into DH5α cells (Invitrogen) according to the manufacturer’s guidelines and grown on LB agar plates containing gentamycin (10 μg/ml) overnight at 37°C. Constructs were validated by colony PCR and sequencing. The ARF sequences within pDONR207 were then recombined into the pGAD-T7 and pGBK-T7 (Clontech) expression vectors by LR clonase reactions, transformed into E. coli, and once more validated by colony PCR and sequencing. The expression vectors were then co-transformed into AH109 Saccharomyces cerevisiae (Clontech), by lithium acetate mediated transformation (Gietz and Schiestl, 2007). Different combinations of AD (pGAD-T7) and BD (pGBK-T7) vectors, or empty vector controls, were introduced in pairs to competent yeast cells to determine whether ARF4 and ETT interact, and through which domains of ARF4. Transformants were plated onto YSD -W-L media and grown at 28°C for 72 hours, before serial dilution (10^0^, 10^1^, 10^2^, and 10^3^) and plated onto YSD -W-L-H-A plates to assess interactions. After plating on selective media, yeast colonies were photographed after growth at 28°C for 7 days. Figures were created using images from the 10^1^ dilution in all cases. Auxin sensitivity was assessed by adding 100 μM IAA (Sigma) to the medium.

### Confocal Microscopy

Stage 8-14 gynoecia were dissected from the plant and placed onto slides in water containing 0.015% Silwet L-77 (De Sangosse Ltd.) to prevent the formation of bubbles. Cover slips were then applied, and the gynoecia were imaged using a Leica SP5 confocal microscope (HyD detector, Smart Gain 30% 200hz, 4x Line Average) with a 40x water immersion lens. GFP was excited at a wavelength of 488 nm and detected at 498–530 nm. Images were taken as Z-stacks, and then processed in Image J 1.48 to form composite 3D projections, which were then converted to RGB images.

## Supporting information

Supplemental Dataset 1-FPKMs all expressed genes

Supplemental Dataset 2-GO Terms and background list

Supplemental Dataset 3- RNAseq Auxin Independent Analysis

Supplemental Dataset 4- RNAseq Auxin Sensitive Analysis

## Author contribution

H.M.M., Y.D. and L.Ø. devised the experiments. H.M.M. conducted the experiments in Arabidopsis with assistance from T. L. and B. N., Y.D. and T.L. conducted experiments in Capsella. H.M.M. wrote the manuscript. H.M.M., Y.D. and L.Ø revised the manuscript. All authors participated in the discussion of the data and in the production of the final version of the manuscript.

## Acknowledgements

Apologies to any colleagues whose work we have not cited due to space limitations. Special thanks to Henrik Jönsson (SLCU) for supporting the completion of the project. Thanks to the JIC microscopy team and both JIC and SLCU horticulture teams for making this work possible. H.M.M. was funded by the John Innes Foundation (at JIC) and the Gatsby Charitable Foundation (at SLCU, GAT3395-PR4B), B.N. by BBSRC (BB/S002901/1) and L.Ø. by BBSRC (BB/P013511/1). The work in Capsella was supported by grants from the National Natural Science Foundation of China (32170227 and 32221001 to Y.D.) and K.C. Wong Education Foundation (GJTD-2020-05, received by Y.D.).

