## Supplemental Data and Methods for "Conserved auxin signalling synergism through the canonical and ETT-mediated pathways promotes gynoecium development in members of the *Brassicaceae*"

Supplemental Figures

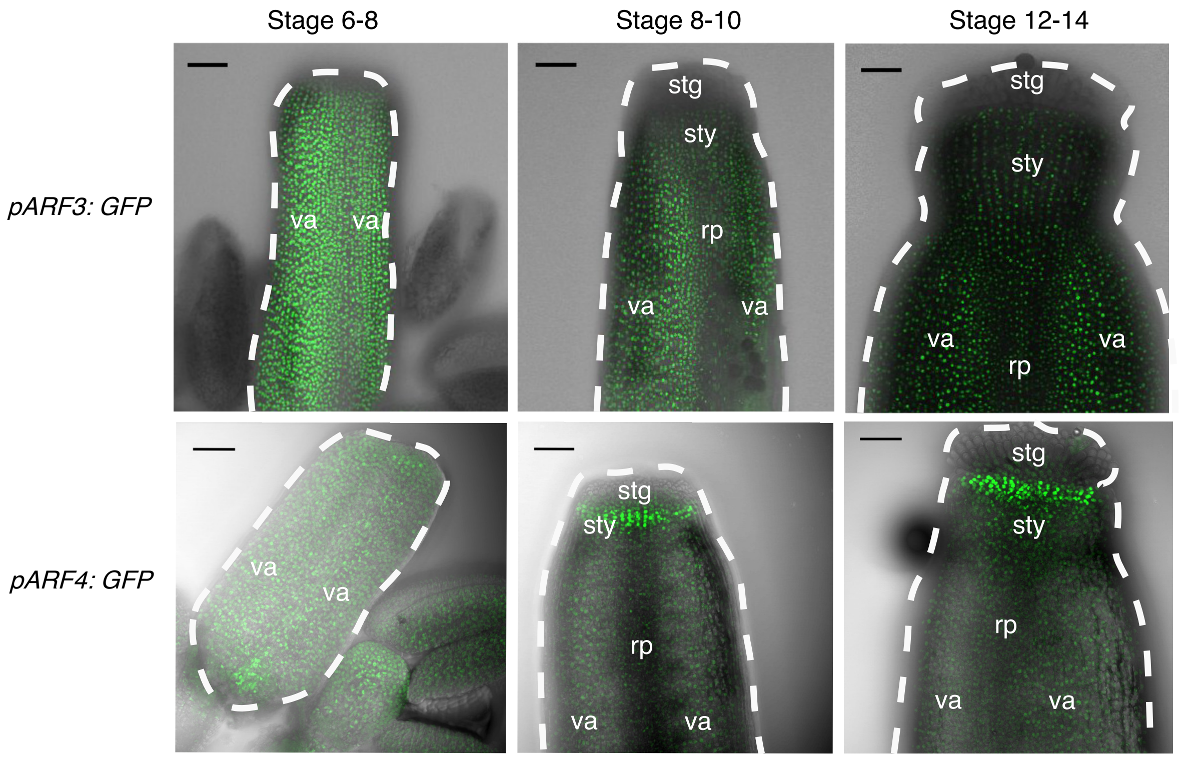


**Fig.S1: Expression of *pETT(ARF3):GFP* and *pARF4: GFP* from stage 6-14 of gynoecium development in *Arabidopsis*.** Outline of gynoeium in the brightfield image is shown by the dashed line. Va, valve, stg, stigma, rp, replum, sty, style. Scale bars 50 μm.

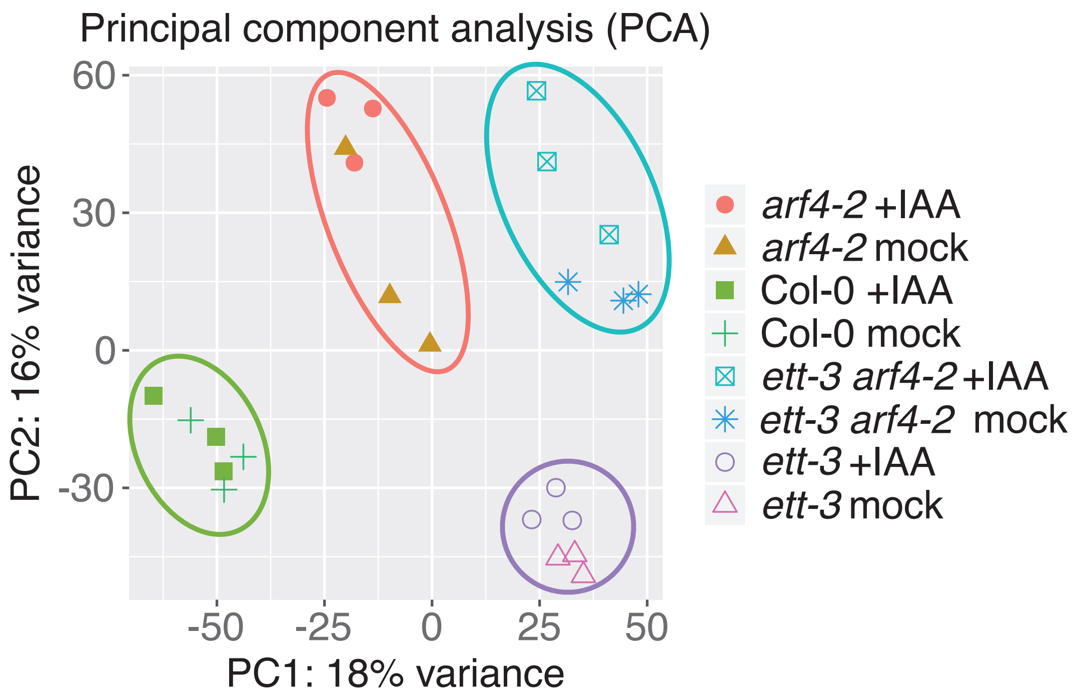


**Fig.S2: Principal component analysis showing clustering of data from each line in the comparative RNA seq experiment in *Arabidopsis*.**

Table S1: Genes which have previously been reported as auxin sensitive by Paponov *et al*., 2008 which were also found to be differentially expressed in response to auxin in Col-0 in the RNA-sequencing analysis.

| Gene | ATG Number | Log2Fold Value +IAA | p Value | Significance | Process |
|---|---|---|---|---|---|
| *GH3.3* | *AT2G23170* | 2.43 | 1.84E-61 | TRUE | Auxin Response |
| *IAA16* | *AT3G04730* | 0.41 | 1.15E-06 | TRUE | Auxin Signalling |
| *IAA2* | *AT3G23030* | 1.08 | 1.49E-11 | TRUE | Auxin Signalling |
| *IAA4* | *AT5G43700* | 0.94 | 3.46E-04 | TRUE | Auxin Signalling |
| *SAUR21* | *AT5G18030* | 1.42 | 8.02E-07 | TRUE | Auxin Response |
| *SAUR50* | *AT4G34760* | 0.55 | 1.51E-05 | TRUE | Auxin Response |
| *SAUR64* | *AT1G29450* | 1.35 | 1.38E-07 | TRUE | Auxin Response |
| *SAUR66* | *AT1G29500* | 0.74 | 3.99E-06 | TRUE | Auxin Response |
| *SAUR67* | *AT1G29510* | 0.91 | 2.75E-09 | TRUE | Auxin Response |
| *PIN3* | *AT1G70940* | 0.43 | 6.97E-05 | TRUE | Auxin Transport |
| *PIN4* | *AT2G01420* | 0.68 | 2.36E-07 | TRUE | Auxin Transport |
| *ARF10* | *AT2G28350* | 0.50 | 1.23E-05 | TRUE | Auxin Signalling |
| *CYP79B2* | *AT4G39950* | 0.60 | 1.45E-03 | TRUE | Auxin Biosynthesis |

| | | | | | |
|---|---|---|---|---|---|
| *ARR16* | *AT2G40670* | -0.48 | 9.95E-04 | TRUE | Cytokinin Signalling |
| *CKX6* | *AT3G63440* | 0.73 | 1.53E-05 | TRUE | Cytokinin Degradation |
| *AT5G43450* | *AT5G43450* | 1.54 | 6.68E-06 | TRUE | Ethylene Biosynthesis |
| *ERF104* | *AT5G61600* | -0.60 | 0.00134427 | TRUE | Ethylene Signalling |
| *ERF113* | *AT5G13330* | -0.52 | 0.00023729 | TRUE | Ethylene Signalling |
| *ERF114* | *AT5G61890* | -1.54 | 6.58E-07 | TRUE | Ethylene Signalling |
| *GA20OX2* | *AT5G51810* | 0.70 | 6.46E-05 | TRUE | GA Biosynthesis |

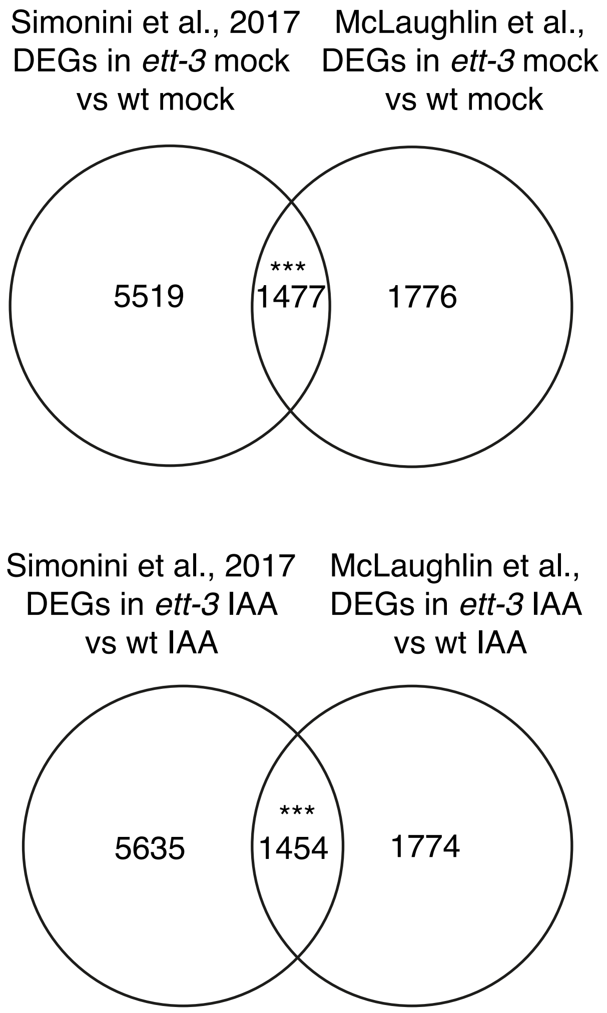


**Fig.S3 Venn diagrams summarizing the overlap between the RNA- sequencing data from Simonini et al. 2017 and those presented in this study.** DEG, differentially expressed genes. Statistical analysis of overlap was conducted using Fisher's Exact Test $p<0.05$. ***= $p<0.001$

Table S2 Statistical assessment of overlap between the Simonini *et al*. 2017 and McLaughlin 2023 datasets (DEG in *ett-3* relative to wildtype after mock treatment). Statistical analysis was conducted using Fisher's Exact Test ($p<0.05$).

| **Analysis parameter** | **value** |
|---|---|
| Overlapping DEGs | 1477 |
| p value (Fisher's Exact Test) | $2.8 x 10^{-146}$ |
| Jaccard Index | 0.168 |
| Significance | TRUE |
| odds ratio | 2.746 |

Table S3: Statistical assessment of overlap between the Simonini *et al*. 2017 and McLaughlin 2023 datasets (DEG in *ett-3* relative to wildtype after IAA treatment). Statistical analysis was conducted using Fisher's Exact Test ($p<0.05$).

| **Analysis parameter** | **value** |
|---|---|
| Overlapping DEGs | 1454 |
| p value (Fisher's Exact Test) | $2.4 x 10^{-134}$ |
| Jaccard Index | 0.164 |
| Significance | TRUE |
| odds ratio | 2.638 |

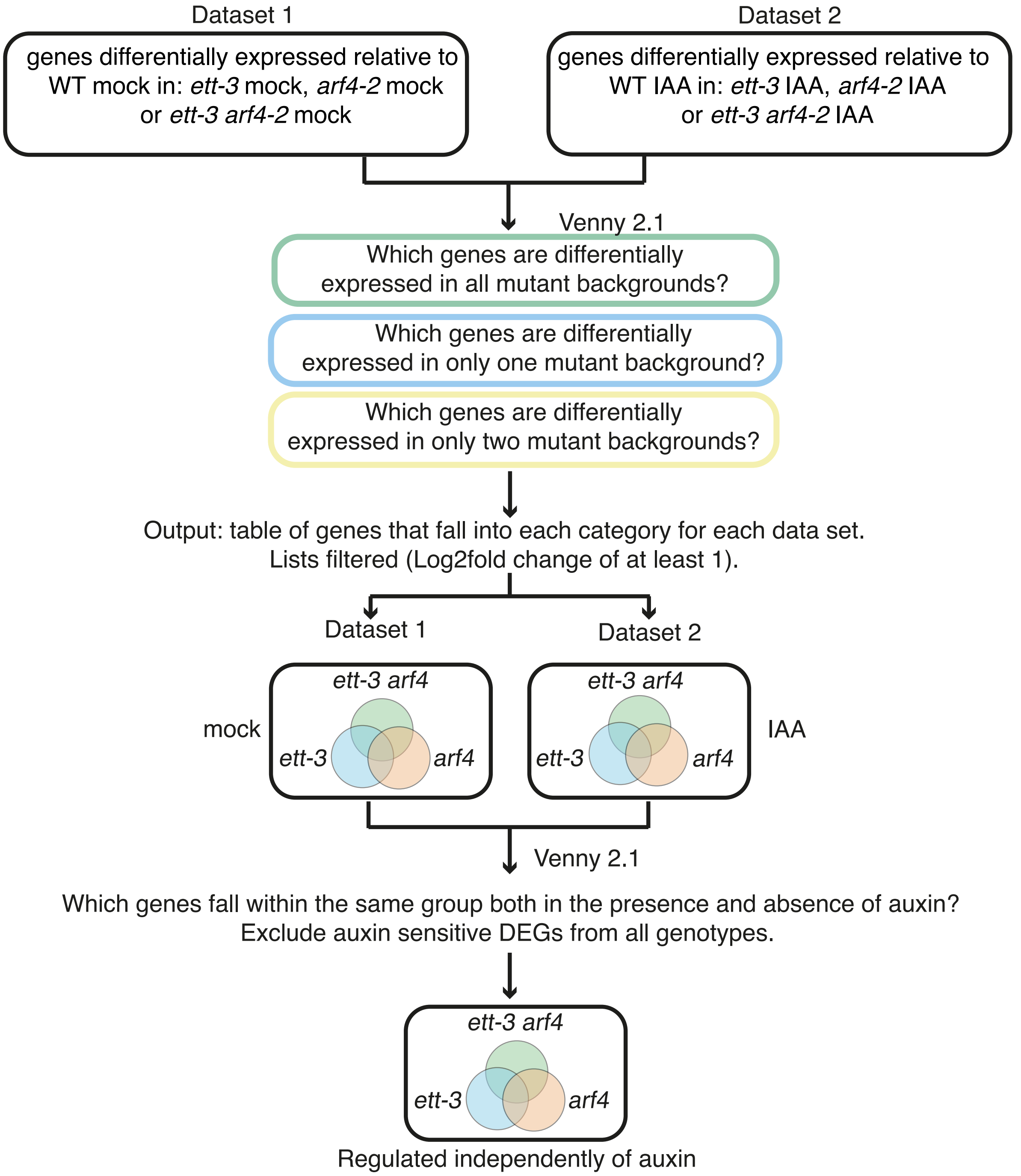


Fig.S4: Schema of auxin independent RNA sequencing analysis. DEG, differentially expressed gene. Two datasets (containing three lists of genes each) were used for this analysis. The first dataset contained lists of genes which were differentially expressed relative to wildtype in *ett-3*, arf4-2 and *ett-3 arf4-2* after mock treatment, whilst the second comprised the same after IAA treatment. The lists were compared in Venny 2.1 to generate two three-way Venn diagrams. Comparisons between the two datasets identified genes which are mis-regulated in the same manner after both mock and auxin treatment. Auxin sensitive genes were removed, leaving genes which are regulated by ETT and ARF4 independently of auxin.

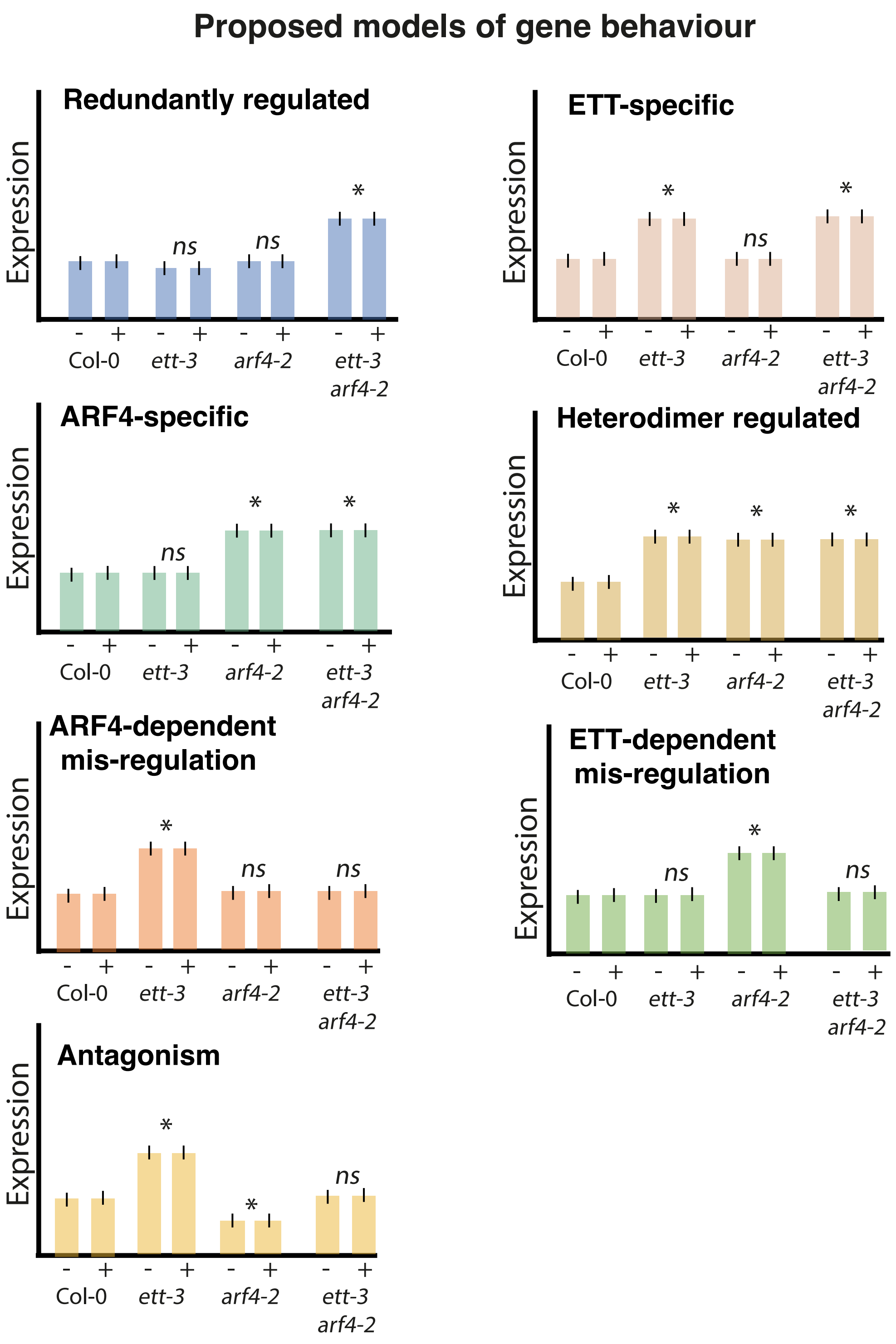


**Figure S5: Proposed models of gene behaviour in each group identified in the auxin sensitive analysis.* denotes the pattern of significance expected in each group, whilst the genes in the example are up-regulated, they could equally show down-regulation.**

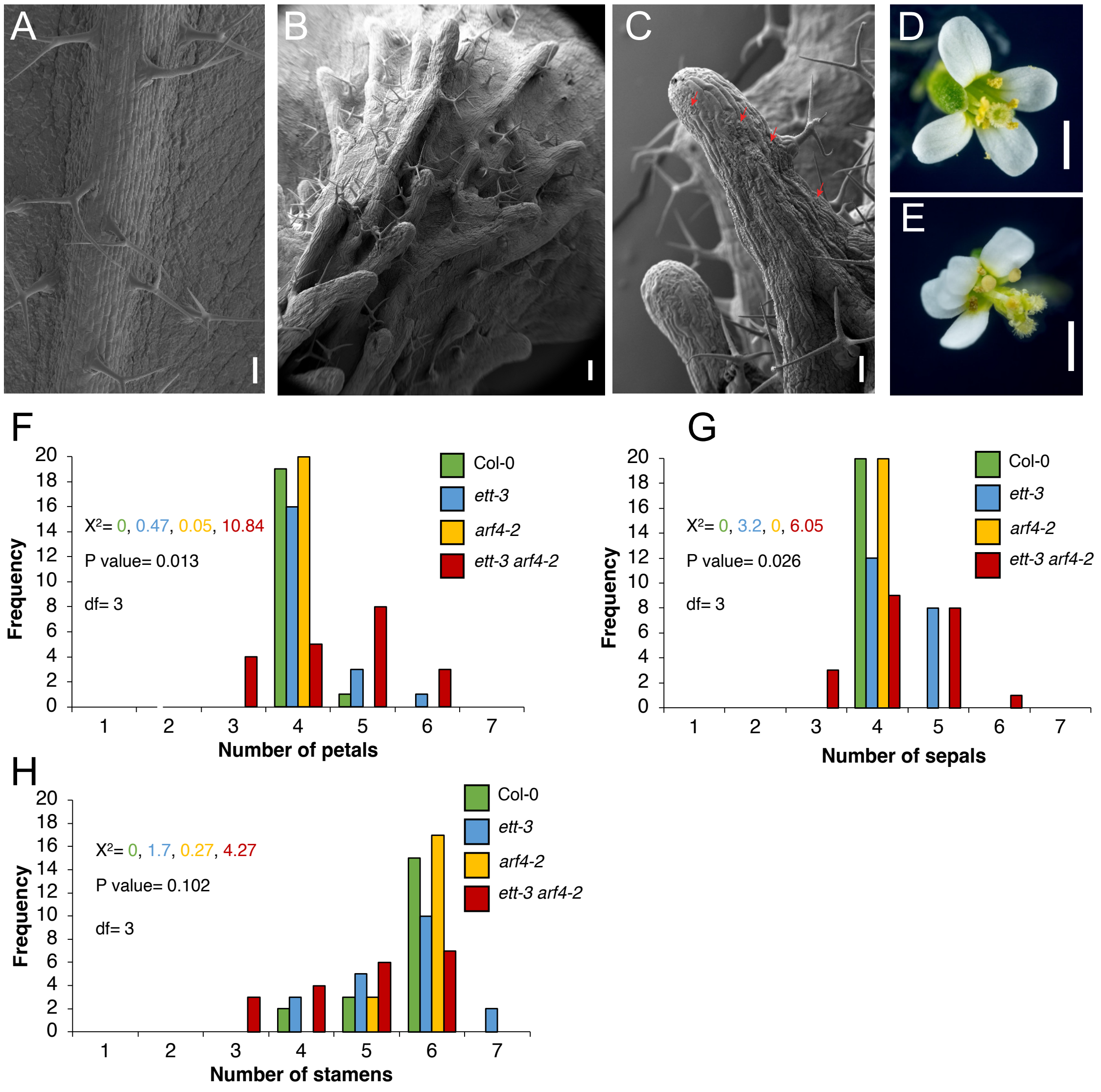


**Fig. S6: Phenotypes in mutant lines correlate with the genes mis-regulated in the auxin independent analysis.** Scanning electron micrographs of the abaxial surface of: A) Col-0 leaf and midvein, B) and C) *ett-3 arf4-2* leaf and blade like projections formed from midvein-like structures. The red arrows in C show stomata developing on the mid-vein like projections of *ett-3 arf4-2* mutants. Photographs of flowers from Col-0 D), and *ett-3 arf4-2* E). Graphs showing variable floral organ number phenotypes observed for petals F), sepals G), and stamens H) in *ett-3* and *ett-3 arf4-2* mutants. In floral organ count studies, n=20 for all lines. Scale bars A) and B) 200 μm, C) 100 μm, D and E) 1cm.

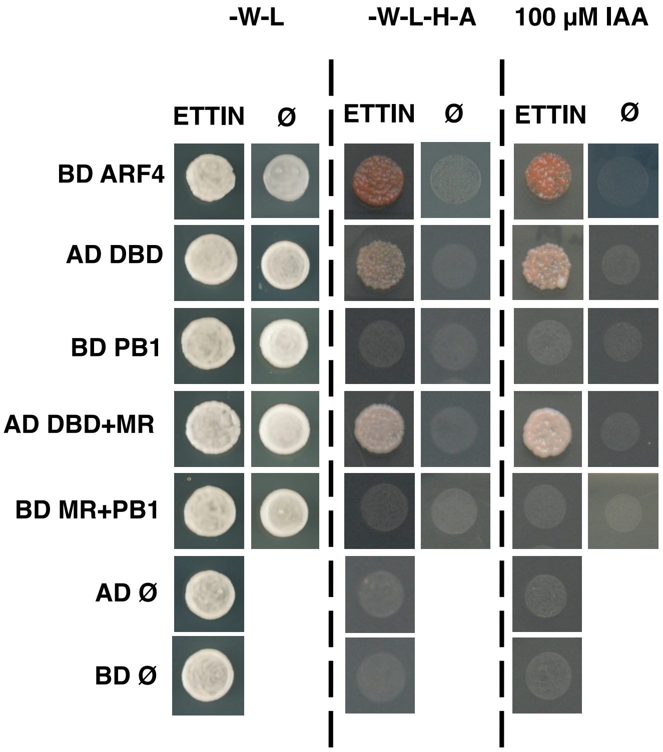


**Fig.S7:** Yeast 2 hybrid assay assessing the interactions between ETT and ARF4. The assay tested interactions of full length ETT with full length ARF4, the ARF4 DNA binding Domain (DBD, amino acid 0-308), ARF4 PB1 domain (PB1, amino acid 663-788), the ARF4 DBD and middle region (MR, amino acid 1-663), and the ARF4 MR and PB1 domain (amino acid 308-788), and empty vector controls (ø).

-W -L media lacking tryptophan and leucine, -W-L-H-A selection media lacking tryptophan, leucine, histidine, and alanine. Auxin treatment was conducted on the -W-L-H-A media.

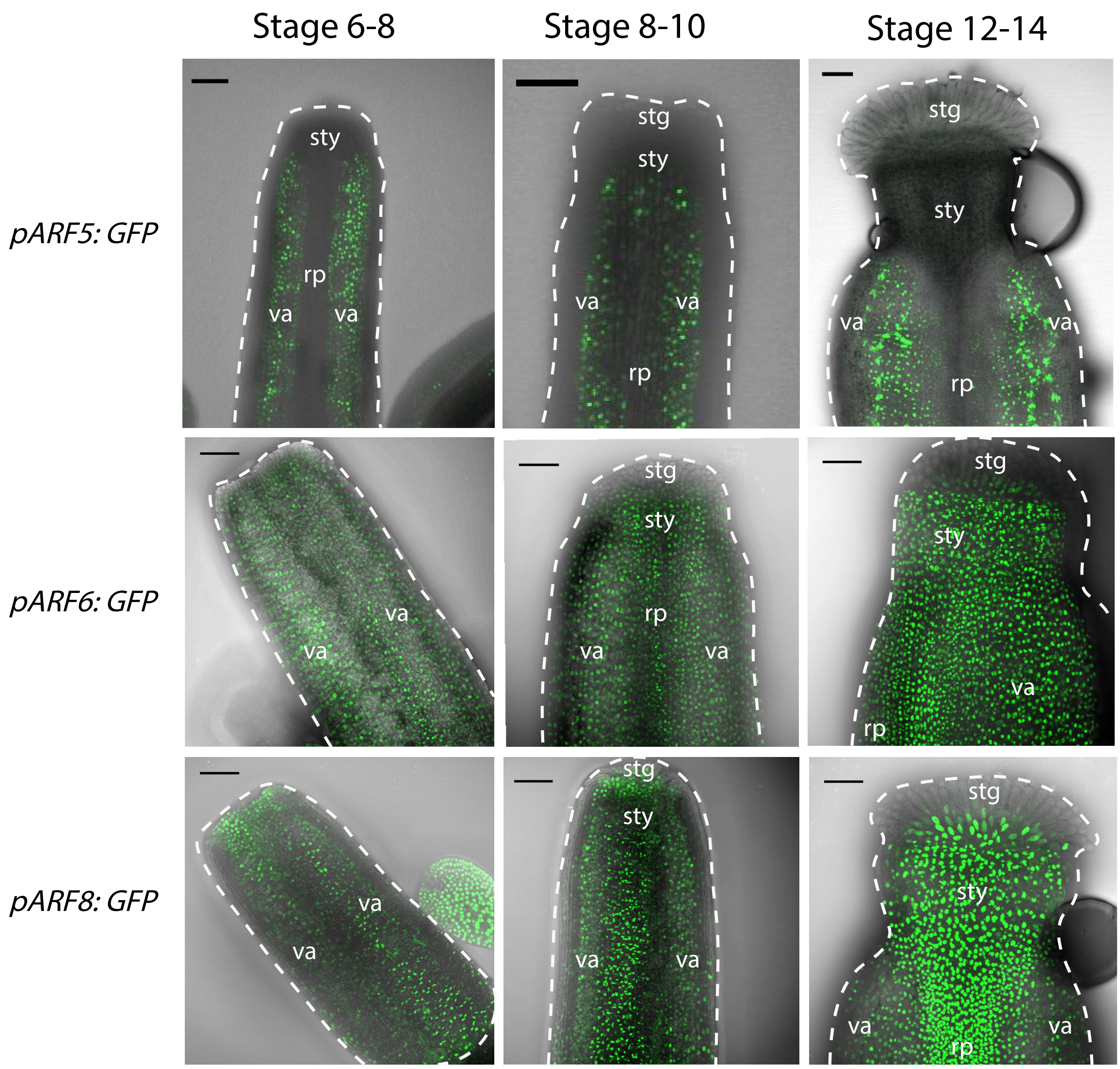


**Expression of *pARF5:GFP*, *pARF6:GFP and pARF8:GFP* from stage 6-14 of gynoecium development in *Arabidopsis*.** Outline of gynoeium in the brightfield image is shown by the dashed line. Va, valve, stg, stigma, rp, replum, sty, style. Scale bars 50 µm.

Table S4: Information on primers, band sizes and optimal $T^a$ for each reaction in Arabidopsis line genotyping.

| Locus | Type | primers | Genotyping by | Size of product (bp) | Optimal $T^a$ (°C) |
|---|---|---|---|---|---|
| *tir1-1* | EMS | HM86+87 | BsaI digestion | WT: 517<br>Mutant: 437 and 80 | 58 |
| *ett-3* | EMS | HM39+40 | sequencing | 246 | 46 |
| *arf4-2* | SALK | WT:HM27+28,<br>Mutant:HM28+LBb1.3 | SALK PCRs | WT: 1196<br>Mutant: 577-877 | 58<br>58 |
| *afb2-3* | SALK | WT: HM31+32<br>Mutant: HM32+LBb1.3 | SALK PCRs | WT: 1131<br>Mutant:598-898 | 56<br>58 |

Supplemental Table S5: Primer sequences used for genotyping Arabidopsis lines and generation of Y2H constructs in this study

| Primer | Sequence | Used for |
|---|---|---|
| HM86 | GCGAATAGCCTTGTCGTTTCC | genotyping |
| HM87 | TCCTAGAGACACAAGGGAGTCTT | genotyping |
| HM39 | CAGATCCCCTGGGATTATAAGTGG | genotyping |
| HM40 | AGACACGCTGTTGATGCCCATTTG | genotyping |
| HM27 | GTGCGTCCTCTCATCTTTTTG | genotyping |
| HM28 | CGCGACAAATCAATAGCTCTC | genotyping |
| HM31 | TCAACGGTCAAGATCCATCTC | genotyping |
| HM32 | CTGCAATTAGCGGCAATAGAG | genotyping |
| LBb1.3 (T-DNA) | ATTTTGCCGATTTCGGAAC | genotyping |
| attB1_ARF4_PB1_F1 | GGGGACAAGTTTGTACAAAAAAGCAGGCTCCAAAAGAATCTGTACAAAGGTT | Amplifying PB1 domain ARF4 Y2H |

| | | |
|---|---|---|
| attB2_ARF4_DBD_1R | GGGGACCACTTTGTACAAGAAAGCTGGGT CTGTAGATACAGCATTAGCCAC | Amplifying DBD ARF4 Y2H |
| attB2_ARF4_MR_R1 | GGGGACCACTTTGTACAAGAAAGCTGGGT CGCTCGAGCTTTGCGGCTTAG | Amplifying ARF4 DBD and MR Y2H |
| attB1_ARF4_MR_PB1_1F | GGGGACAAGTTTGTACAAAAAAGCAGGCT CC GCCAAGAAATGGACTTCCTGACTCA | Amplifying ARF4 MR and PB1 domain Y2H |
| attB2_ARF4_MR_PB1_1R | GGGGACCACTTTGTACAAGAAAGCTGGGT C TCAAACCCTAGTGATTGTAGGAGAAG | Amplifying ARF4 MR and PB1 domain Y2H |
| attB1_ETT_1F | GGGGACAAGTTTGTACAAAAAAGCAGGCT CCATGGGTGGTTTAATCGATCTG | Amplifying Full Length ETT Y2H |
| attB2_ETT_1R | GGGGACCACTTTGTACAAGAAAGCTGGGT CCTAGAGAGCAATGTCTAGCAACA | Amplifying Full Length ETT Y2H |
| attB1_ARF4_1F | GGGGACAAGTTTGTACAAAAAAGCAGGCT CCATGGAATTTGACTTGAATACT | Amplifying Full Length ARF4 Y2H |
| attB2_ARF4_1R | GGGGACCACTTTGTACAAGAAAGCTGGGT C TCAAACCCTAGTGATTGTAG | Amplifying Full Length ARF4 Y2H |

Table S6: List of primers used for the construction of GUS reporter lines, genome editing and genotyping in *Capsella*.

| Primer | Sequence (5’ to 3’) | Used in experiment |
|---|---|---|
| *pCrAFB2-F* | *CCTCTAGAGTCGACCTGCAGTCAA CCTACTGACTACGTATC | GUS line construction |
| *pCrAFB2-R* | *CTCAGATCTACCATGGCTCTCCAG TAACTTGAACAAC | GUS line construction |
| *pCrAFB3-F* | *CCTCTAGAGTCGACCTGCAGGAG TTTTATTATGAAGCCTTT | GUS line construction |
| *pCrAFB3-R* | *CTCAGATCTACCATGGCTTCCGAA ACCCAAACAATAGG | GUS line construction |
| *pCrTIR1-F* | *CCTCTAGAGTCGACCTGCAGTGAC CTCTCTAATCGCATCC | GUS line construction |

| | | |
|---|---|---|
| *pCrTIR1-R* | *CTCAGATCTACCATGGTGCGCCCA AGAATAACTTCG | GUS line construction |
| *pCrETT-F* | *CCTCTAGAGTCGACCTGCAGAAA ACGCACCAAGTAATTATGA | GUS line construction |
| *pCrETT-R* | *CTCAGATCTACCATGGTATAGAG AGAGAGACAGAGATA | GUS line construction |
| *pCrARF4-F* | *CCTCTAGAGTCGACCTGCAG GTTTGGTTTGGCCATGTATGAC | GUS line construction |
| *pCrARF4-R* | *CTCAGATCTACCATGGTGAAAAA AAAAAAGCTTCTTAGAGGA | GUS line construction |
| *pAtAFB2-F* | CCTCTAGAGTCGACCTGCAGATTG TCTTGAATAATTGTTTCATCCATA | GUS line construction |
| *pAtAFB2-R* | *CTCAGATCTACCATGGCTCTAACT TCACCAGCAAGATTCC | GUS line construction |
| *pAtAFB3-F* | *CCTCTAGAGTCGACCTGCAGCAC ATGGCAGCCATTGTAGC | GUS line construction |
| *pAtAFB3-R* | *CTCAGATCTACCATGGCTTCCTCT ATTGATTGTCGAACAGAG | GUS line construction |
| *pAtTIR1-F* | *CCTCTAGAGTCGACCTGCAGTCAT TCAGAGTTGGATCTAATATCGCT | GUS line construction |
| *pAtTIR1-R* | *CTCAGATCTACCATGGTGCGGCCA AATAACCTCG | GUS line construction |
| *CrAFB2-Exon2-gRNA1* | CTTTAGAGAGGCTTGTTGCA | Gene editing |
| *CrAFB3-Exon1-gRNA1* | GAACTGCTACGCCATCAACC | Gene editing |
| *CrTIR1-Exon1-gRNA1* | ATTAGCCCTGTCGTTCCCGG | Gene editing |
| *CrARF4-Exon1-gRNA1* | GTAGTTGTGTATTTCCCACA | Gene editing |
| *CrETT-Exon1-gRNA1* | TCTGAACGTGATGGAGACGG | Gene editing |
| *CrAFB2-Geno-F* | GCATCTTCGGGATCTGGACTTG | Gene CRISPR genotyping |
| *CrAFB2-Geno-R* | TTTCTTTATGGCAGCCTTGAGT | Gene CRISPR genotyping |
| *CrAFB3-Geno-F* | TCATCGCTTCTCACAGAGACA | Gene CRISPR genotyping |
| *CrAFB3-Geno-R* | AATGGAAGCTAAGCCGTCAG | Gene CRISPR genotyping |
| *CrTIR1-Geno-F* | TTCGGACTTCTTTCCCCAACTCTG | Gene CRISPR genotyping |
| *CrTIR1-Geno-R* | TCAGCGAAGTGAGGTTTCCCCT | Gene CRISPR genotyping |

| | | |
|---|---|---|
| *CrETT-Geno-F* | TGCTAGTCTCTCTTCTTTTGGCT | Gene CRISPR genotyping |
| *CrETT-Geno-R* | AACAAAACAAACCCCCATAC | Gene CRISPR genotyping |
| *CrARF4-Geno-F* | CCTCTTCCACAGGCTCTGCCTC | Gene CRISPR genotyping |
| *CrARF4-Geno-R* | AGGCGTTCGTTTAACGGTTGAG | Gene CRISPR genotyping |

*The sequences shaded in yellow indicate the homologs to the plasmid and used for In-Fusion cloning.

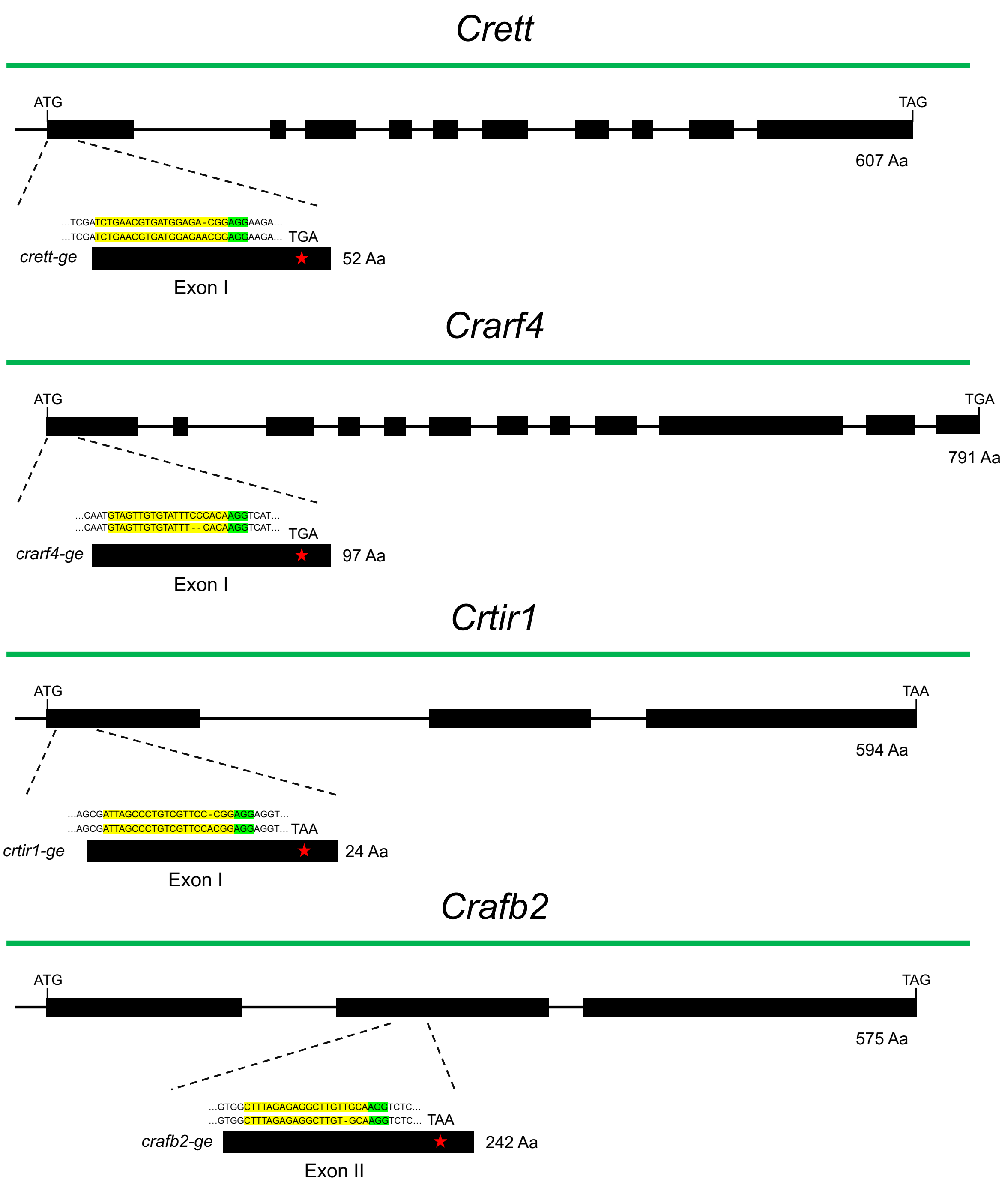


Fig.S9. Information on the mutations induced by CRISPR/Cas9 in *Capsella rubella*. In *Crett*, *Crarf4*, and *Crtir1*, early stop codons were induced by deletions in exon one of the genomic sequence. In *Crafb2*, an early stop codon was produced in exon two. The guide RNAs and PAM sequences were highlighted with yellow and green, respectively.
